# High resolution cryo EM analysis of HPV16 identifies minor structural protein L2 and describes capsid flexibility

**DOI:** 10.1101/2020.10.27.357004

**Authors:** Daniel J. Goetschius, Samantha R. Hartmann, Suriyasri Subramanian, Carol Bator, Neil D. Christensen, Susan L Hafenstein

## Abstract

Human papillomavirus (HPV) is a significant health burden and leading cause of virus-induced cancers. HPV is epitheliotropic and its replication is tightly associated with terminal keratinocyte differentiation making production and purification of high titer virus preparations for research problematic, therefore alternative HPV production methods have been developed for virological and structural studies. In this study we use HPV16 quasivirus, composed of HPV16 L1/L2 capsid proteins with a packaged cottontail rabbit papillomavirus genome. We have achieved the first high resolution, 3.1Å, structure of HPV16 by using a local subvolume refinement approach. The high resolution enabled us to build L1 unambiguously and identify L2 protein strands. The L2 density is incorporated adjacent to conserved L1 residues on the interior of the capsid. Further interpretation with our own software for Icosahedral Subvolume Extraction and Correlated Classification (ISECC) revealed flexibility, on the whole-particle level through diameter analysis and local movement with inter-capsomer analysis. Inter-capsomer expansion or contraction, governed by the connecting arms, showed no bias in the magnitude or direction of capsomer movement. We propose that papillomavirus capsids are dynamic and capsomers move as rigid bodies connected by flexible linkers. The resulting virus structure will provide a framework for continuing biochemical, genetic and biophysical research for papillomaviruses. Furthermore, our approach has allowed insight into the resolution barrier that has previously been a limitation in papillomavirus structural studies.

## Introduction

Human papillomaviruses (HPVs) cause epithelial tumors and are the etiologic agents of numerous anogenital and oropharyngeal cancers(Bosch et al., 1995; Crow, 2012; Walboomers et al., 1999). HPV is implicated in more than 70% of head, neck, and throat cancers, the occurrence of which has been increasing steadily, specifically in a younger group of non-smoking 40-50 year old white men (Lewis et al., 2015; Lydiatt et al., 2017). Although there are over one hundred known genotypes, HPV16 is associated with 50% of all cervical cancers and 90% of all other HPV-containing anogenital and oropharyngeal malignancies(zur Hausen, 1991). We used HPV16 in this study due to its disproportionate contribution to human disease.

HPV is epitheliotropic and its replication is tightly associated with terminal keratinocyte differentiation making production and purification of high-titer virus preparations for research problematic. Consequently, alternative HPV production methods of virus-like particles (VLPs) have been developed within the HPV community for virological and structural studies. Capsid protein only VLPs can be composed of the major structural protein (L1) alone, or of the major and minor capsid proteins (L1/L2) and are not infectious since they are devoid of viral genomes(Hernandez et al., 2012; Kirnbauer et al., 1992, 1993). Pseudovirus is comprised of both structural proteins (L1/L2) with plasmid DNA packaged as a mock genome(Buck, Pastrana, et al., 2005; Buck, Thompson, et al., 2005). Quasivirus is similarly comprised of both structural proteins (L1/L2), but packaged with a cottontail rabbit papillomavirus genome (CRPV) to assemble a structurally complete L1/L2 capsid that is infectious (Christensen, 2005). All these different HPV VLPs preserve the main attributes of the native capsid structure and have been used successfully for vaccine development and for studies of antigenicity, receptor usage, entry mechanisms, and structural analyses. Quasiviruses were used throughout the work described here.

Human papillomaviruses are non-enveloped, circular dsDNA containing viruses with capsids comprised of 360 copies of the major capsid protein, L1, and an uncertain number of the minor structural protein, L2(Buck et al., 2008a; Finch & Klug, 1965). Five copies of L1 intertwine to form each of the 72 capsomers that make up the *T*=7*d* icosahedron. The L2 structure is unknown and its capsid incorporation is unclear. L1 takes the form of the ubiquitous jellyroll of anti-parallel beta strands (BIDG and CHEF), which are connected by flexible loops that extend outward from the surface of the capsomer and constitute the majority of the L1 hypervariable regions. Pentavalent capsomers lie on the icosahedral five-fold axis and are surrounded by five neighboring capsomers. The remaining sixty hexavalent capsomers are each bordered by six capsomers. To form the icosahedral capsid the pentavalent and hexavalent environments confer quasi-equivalent conformations of L1. The asymmetric unit is comprised of six L1s, five of which make up a single hexavalent capsomer and one L1 that contributes to a pentavalent capsomer (Fig 1.A-B). An extension of each L1 C-terminal arm links capsomers through the formation of disulfide bonds between Cys 428 and Cys 175 of the adjacent capsomer (Buck, Thompson, et al., 2005; Sapp et al., 1998).

**Figure 1.**
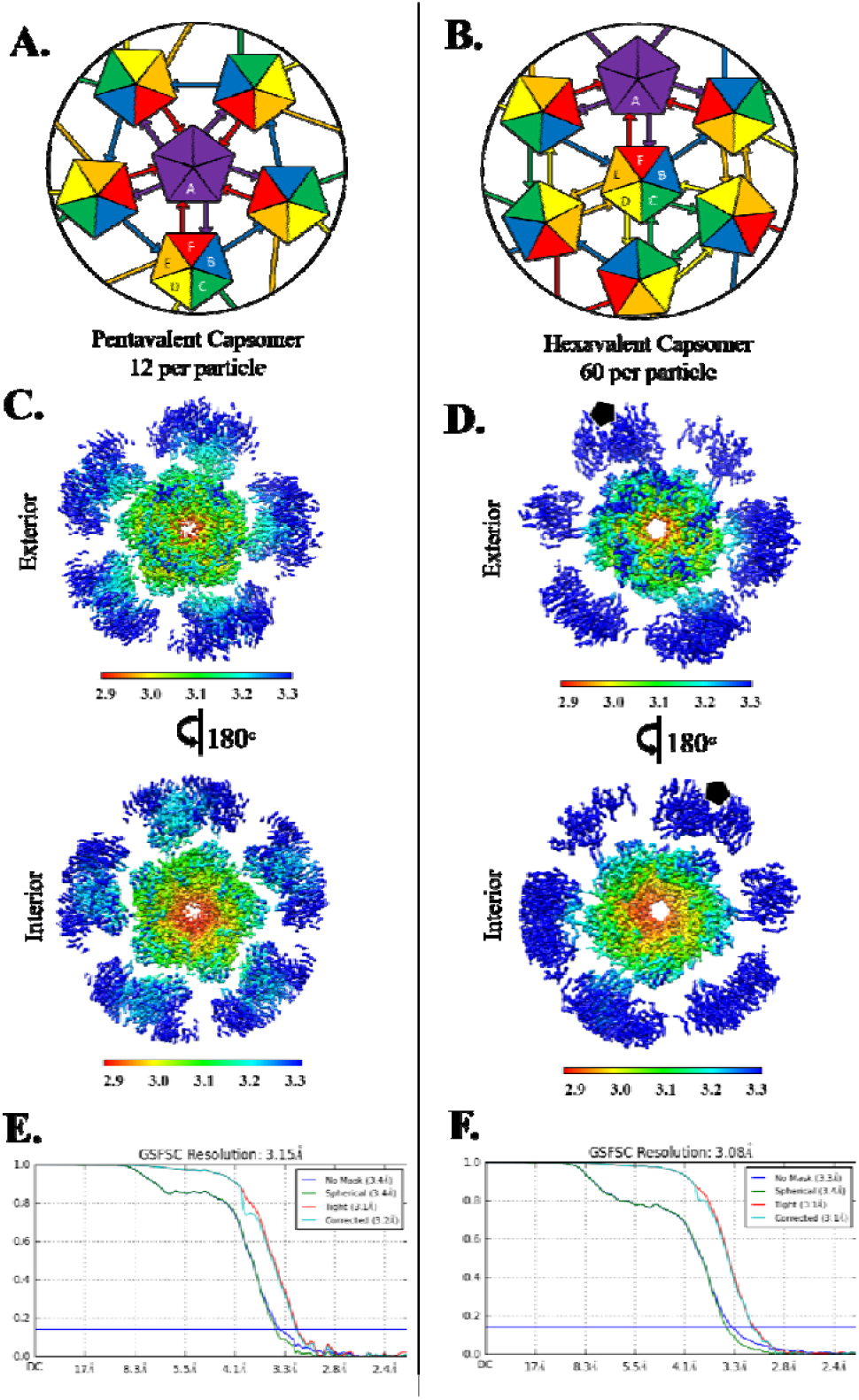
Capsomer environments and subparticle refinement. (A, B) Illustrated graphically by cartoons of pentavalent and hexavalent capsomers, the HPV capsid asymmetric unit is made up of six L1 chains, labeled chain A-F and colored purple, blue, green, yellow, orange, and red respectively. (A) Each A chain of the pentavalent capsomer makes a connection with chain F of the neighboring hexavalent capsomer, showing fivefold icosahedral symmetry. The pentavalent capsomer is surrounded by single arm connections of chain B to chain E of hexavalent capsomers in a counterclockwise ring contribution. (B) Each C chain (green) can be seen interacting with another C chain, showing threefold icosahedral symmetry. Each E chain (orange) contributes to D chain (yellow), showing twofold symmetry. (C, D) Surface rendered subvolumes colored according to local resolution (color key) with exterior views (upper panels) and views flipped 180° for interior view (lower panel). Highest resolution (red) in both environments is seen at the interior core of the capsomer with resolution diminishing at the connecting arms and at the outer edges (C) Pentavalent subvolumes are centered on the fivefold icosahedral symmetry axes, whereas a black pentagon identifies the fivefold axis location in the hexavalent capsomer (D). (E) Pentavalent capsomer FSC curve to 3.15Å. (F) Hexavalent capsomer FSC curve to 3.08Å.

Previously solved structures of HPV16 capsids have been limited by modest resolution. Although there are high resolution x-ray crystallography maps of the HPV16 L1 pentamer, the complete HPV *T=7d* capsid has only been visualized by cryo EM (X. S. Chen et al., 2000; Dasgupta et al., 2011). Currently, all reconstructions of the entire capsid have been resolved with the use of icosahedral symmetry averaging, and the highest resolution structure is 4.3 Å(Baker et al., 1991; Buck et al., 2008b; Cardone et al., 2014; Guan, Bywaters, Brendle, Lee, Ashley, Makhov, et al., 2015; Guan, Bywaters, Brendle, Lee, Ashley, Christensen, et al., 2015; Guan et al., 2017b; Lee et al., 2015, p. 5). The commonality between all HPV structures that have been published is the reliable fit of only the major capsid protein, L1.

Although controversy remains about the number and positioning of L2 proteins incorporated into the capsid, L2 has been predicted to bind in the center of the capsomer with part of the N-terminus exposed on the surface(Lowe et al., 2008). Some stoichiometric studies have indicated that there are between 12-36 L2 molecules per HPV capsid(Okun et al., 2001). However, other studies have suggested the amount of L2 per capsid is variable, and may include up to 72 L2 molecules per capsid, perhaps one within each capsomer(Buck et al., 2008a; Doorbar & Gallimore, 1987; Trus et al., 1997). Specific L2 functions have been determined and include facilitating the encapsidation of DNA, involvement in the conformational changes of the capsid during entry, disruption of the endosomal membrane, and subcellular trafficking of the viral genome(Bronnimann et al., 2013; Buck et al., 2008b, p. 2; Raff et al., 2013).

There have been many advances in cryo EM mainly in hardware and reconstruction software that have made it possible to obtain atomic resolution structures of viruses and other macromolecules(Bai et al., 2015; Goetschius, Parrish, et al., 2019). Cryo EM reconstructions of icosahedral viruses traditionally use symmetry averaging to refine the capsid; however, more recent software developments including localized reconstruction and block-based reconstruction allow subvolumes to be designated for additional refinement(Ilca et al., 2015), (Zhu et al., 2018). These approaches improve map resolution by reconstructing smaller subvolumes of the virus, which often compensates for capsid-wide flexibility and the defocus gradient over the capsid. These advances have also allowed for resolution of asymmetric features, such as minor capsid proteins that do not follow strict icosahedral symmetry(Goetschius, Parrish, et al., 2019; Ilca et al., 2015; Zhu et al., 2018).

In this study we used a subparticle refinement approach that allowed us to overcome the previous resolution barriers in HPV structural studies. We developed custom software, Icosahedral Subvolume Extraction & Correlated Classification (ISECC), to assess capsid flexibility on a per particle basis. Our high resolution map provided the most reliable structure of L1 to date, including corrections to the previous model. The structure also revealed for the first time unambiguous L2 density adjacent to a conserved L1 loop. Assessment of capsid flexibility revealed dynamic capsids with imperfect icosahedral symmetry. This continuous heterogeneity is the likely cause of limited resolution of HPV structure in previous studies and may play a role in crucial biological processes. The work describes methods to achieve high resolution that will lay the groundwork for future structural studies of papillomaviruses and polyomavirus.

## Results

### Imposing icosahedral symmetry during refinement results in a moderate resolution map

Cryo EM micrographs showed virus particles of approximately 50 nm diameter composed of discrete, discernable capsomers (S.Fig 1.A). Consistent with previous studies, variation existed among virus particles, including the occasional rod-like structure(Guan et al., 2017b). Using a standard processing pipeline (Methods) 202,705 particles were classified to select 181,299 particles that contained internal density consistent with packaged genome. Icosahedral symmetry averaging was imposed during refinement to produce a 4.5 Å resolution map. At this resolution α-helices and β-sheets were discernable; however, the new map provided no improvement over previously published structures and a true atomic model could not be built. This result was disappointing considering the high quality micrographs, a large particle number, well-defined 2D classes (S.Fig 1), and advances in software since the 4.3Å HPV16 structure was published by Guan, et al. in 2017(Bai et al., 2015; Goetschius, Lee, et al., 2019; Guan et al., 2017a, p. 4, 2017b).

### Using a subvolume reconstruction approach resulted in high resolution

Icosahedral averaging alone could not compensate for the heterogeneity among the capsids that is likely due to the flexibility between capsomers. To overcome and characterize capsid flexibility we developed Icosahedral Subvolume Extraction & Correlated Classification (ISECC), a suite of programs inspired by the Localized Reconstruction approach (Abrishami et al., 2020; Ilca et al., 2015). ISECC both defines the subparticles for local refinement and provides metadata for subsequent analysis. Here we defined the subparticles for extraction to correspond to pentavalent and hexavalent capsomers. After extraction, local refinement resulted in subvolume maps with local resolution ranging from 2.9-3.3Å with the beta-jellyroll motif of the capsomer cores attaining better resolution than the solvent-exposed variable loops (Fig 1.C-D). The high resolution capsomer maps were recombined into an icosahedral capsid with 3.1Å resolution overall (ISECC_recombine) (Fig 1.E-F), a dramatic improvement over the original 4.5Å map (Fig 2). The recombined map allowed the full asymmetric unit (Fig 2.C) of HPV to be unambiguously built for the first time.

**Figure 2.**
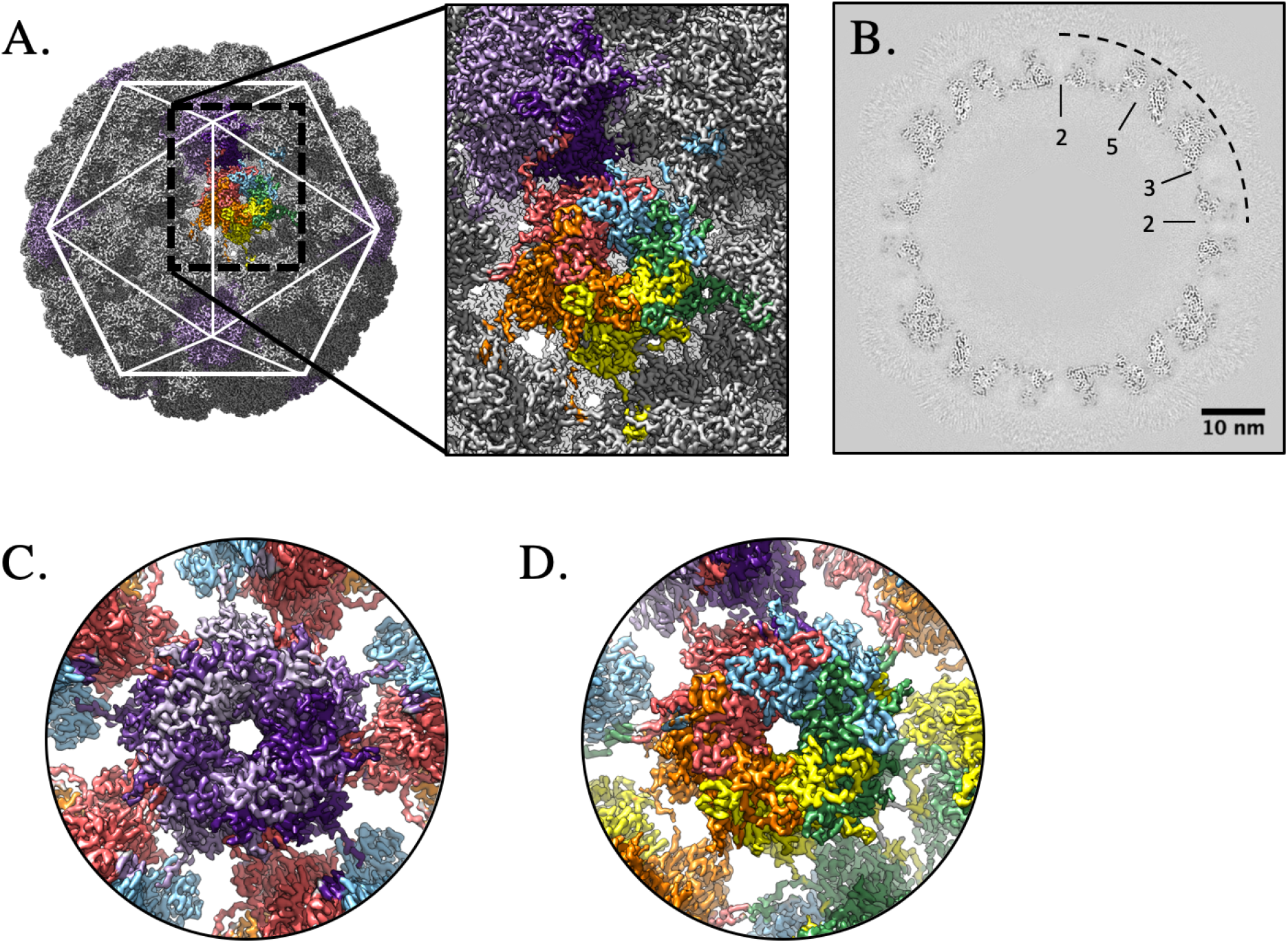
High resolution capsid map. The high resolution subparticle maps were recombined into a complete capsid (gray, surface rendered) (A), allowing the complete asymmetric unit (A, inset, color code as in Fig. 1A) to be visualized in the context of the capsid (icosahedral cage). The high quality of the map is evident in the central section (B) with symmetry axes denoted (black dotted line) and scale bar. Density for individual L1 chains could be continuously traced in both the pentavalent (C) (L1 chain A; shades of purple) and hexavalent (D) capsomer maps (bottom right; L1 chains B-F; blue, green, yellow, orange, red).

Notably, resolution did not improve beyond 3.1Å, despite exceptional subparticle number (pentavalent: 2,175,588, hexavalent 10,877,940), which suggested the presence of additional heterogeneity. However, multiple attempts to 3D classify the subparticles, both before and after local refinement, failed to produce structurally distinct classes. This suggested that the signal-to-noise ratio (SNR) is the limiting factor in achieving truly atomic resolution. Optimized and iterative subtraction of neighboring subparticles might improve the resolution further, but such an experiment would require new software development and achieve modest improvements if successful.

### A new, accurate structure for L1 was built into the high resolution map

The previous L1 structure (PDB ID 5KEP) was accurate to only modest resolution since it was generated from a 4.3 Å cryo EM density map(Guan et al., 2017b). However, as the model contained the asymmetric unit, it was used to initiate the build. Each L1 chain of the asymmetric unit was initially built into the corresponding capsomer subvolume density. The pentavalent capsomer is comprised of chain A with connecting arms from chain F, whereas, the hexavalent capsomer is comprised of chains B-F with the connecting arms from chains A-E (Fig 1A-B).

After refining the structures in each of the subvolumes, the six L1 chains were assembled into the asymmetric unit. This pdb was then validated against the recombined map (S. Table 1). The resolution allowed unambiguous placement of most sidechains (S.Fig 2) and presented continuous density for most of the termini. For the six chains of the asymmetric unit (Fig 1.A-B, chains A-F) there was strong continuous density stretching from Tyr12 to Arg485 (A), Val16 to Phe480 (B), Met1 to Phe480 (C & D), Leu3 to Gly483 (E), and Met1 to Gly483 (F) respectively. The refined structure superimposed with 5KEP with a C-alpha root mean squared deviation (RMSD) of 2.37Å. The improved resolution of our map when compared to the previous 4.3Å structure (Guan *et al.*) allowed us to correct the 36 residues making up the C-terminal arm extensions (402 to 439)(Guan et al., 2017b). This map clearly resolved density corresponding to the Lys475 to Phe480 loop that was previously incorrectly assigned. N-terminal residues starting from Met 1, which had previously been unresolved, were now assigned for most of the chains in the asymmetric unit. Overall, the density in this map allowed clear assignment of residues and corrected the HPV16 L1 structure within the context of the icosahedral capsid.

### Each connecting arm has a different conformation

The pentavalent and hexavalent L1 capsomers share a common core architecture but have different connecting arm structures (residues 402-439). The connecting arm extends from the base of the donor capsomer, looping into the neighboring capsomer to make a disulfide bond between Cys428 of the connecting arm and Cys175 of the neighboring capsomer, and then returns to the donor above the initial extension.

Capsomers superimposed with a c-alpha RMSD of 2.75Å (Fig 3.A); however, after exclusion of connecting arm and N-terminal residues 1-23 (Fig 3.B) the c-alpha RMSD was 0.52Å. These differences are further described by comparing each copy of L1 (chains A-F) that makes up the asymmetric unit (Fig 3.C). The six chains of L1 superimposed with a c-alpha RMSD of 6.16Å, and with the removal of the connecting arms (402-439) and N-terminal residues (1-23) resulted in an RMSD of 0.38Å (Fig 3.D), demonstrating the beta-jellyroll stability in the core of L1. The differing conformations of the connecting arms were then investigated to evaluate the extent of similarities and differences. All six conformations were superimposed from amino acid residues 385 to 472 to include the flanking alpha helices for alignment purposes. The overall RMSD of all six connecting arms was 8.56Å (Fig 3.E). When evaluating the connecting arms pairwise RMSD values ranged from 1.35 to 18.23Å, with the best RMSD values for the pairwise alignment between hexavalent chain F and pentavalent chain A (RMSD:1.35Å) (Fig 3.F).

**Figure 3.**
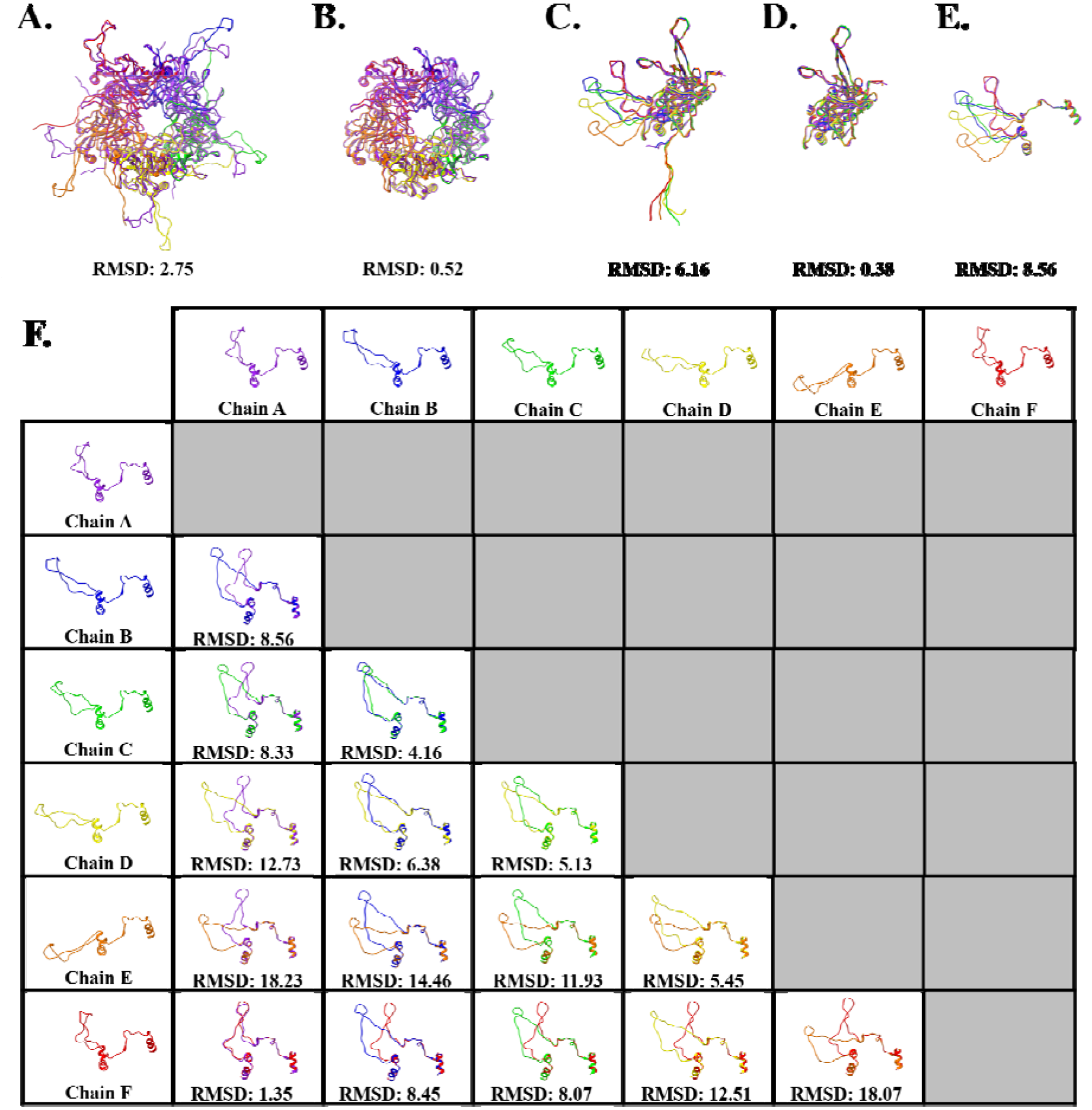
Differences in L1 structural conformation. All RMSD values are given in Angstroms. (A) Pentavalent (five A chains, purple) and hexavalent (chain B-F, blue, green, yellow, orange, and red) capsomers superimposed with an RMSD of 2.75Å. (B) Structure comparison of pentavalent and hexavalent capsomer conformations excluding amino acid side chains 402-439 that extend as a connecting arm to the neighboring capsomer, and the N-terminal residues 1-23. (C) Structure comparison of the six L1 conformations, making up the asymmetric unit chains A-F. (D) Structure comparison of the six L1 conformations, excluding amino acid side chains from the connecting arms and the N-terminal residues. (E) Structure comparison of the connecting arms of the six different L1 conformations found in the asymmetric unit. The flanking secondary structure was included for alignment, depicted as amino acid residues 385-472. (F) Pair-wise structure comparison of the six connecting arm conformations and their associated RMSD values.

### The same flexible region comprised of nine amino acids is seen in each connecting arm conformation

The fit of these connecting arms revealed that the same sequence was found in different conformations. The connecting arms were resolved to a more modest resolution compared to the capsomer core. Specifically, the surface exposed region of the arms from His431 to Asp439 was found to be poorly resolved in each chain, even though the connecting arms are in varying conformations. The most compact conformation was in L1 chains A and F, whereas in chains D and E the conformation was the most extended, with chains B and C exhibiting an intermediate structure (Fig 3.E).

### Both pentavalent and hexavalent capsomers have non-L1 density

After refining L1 into the density of the pentavalent and hexavalent capsomers, there was unfilled density on the interior in two distinct locations (Fig 4.A). These putative L2 densities appeared as strands with protruding knobs consistent with amino acid side chains. The first density, approximately four amino acids, had the overall shape of an arcing fishhook located over L1 Lys475 (Fig 4.B). This fishhook density was present in the previously published structure by Guan *et al.* but was incorrectly interpreted as L1 due to the moderate resolution of the previous map. The second, larger L2 density, approximately six amino acids, was found flanking an L1 loop region that extended from Ser306 to Ile327 (Fig 4.C-D). To assess the region of L1 that flanked this L2 density, conservation of residues within the L1 306-loop was evaluated across the nine types of HPV that are incorporated into the GARDASIL ^®^9 HPV vaccine(Panatto et al., 2015). These clinically relevant types possess >10% sequence diversity in L1; however, the 306-loop was highly conserved and all changes to this region were functionally conserved mutations (Fig 4.E, S. Table 2). Both of these L2 densities were present with different intensities within the pentavalent and hexavalent capsomer environments. (S.Fig 3)

**Figure 4.**
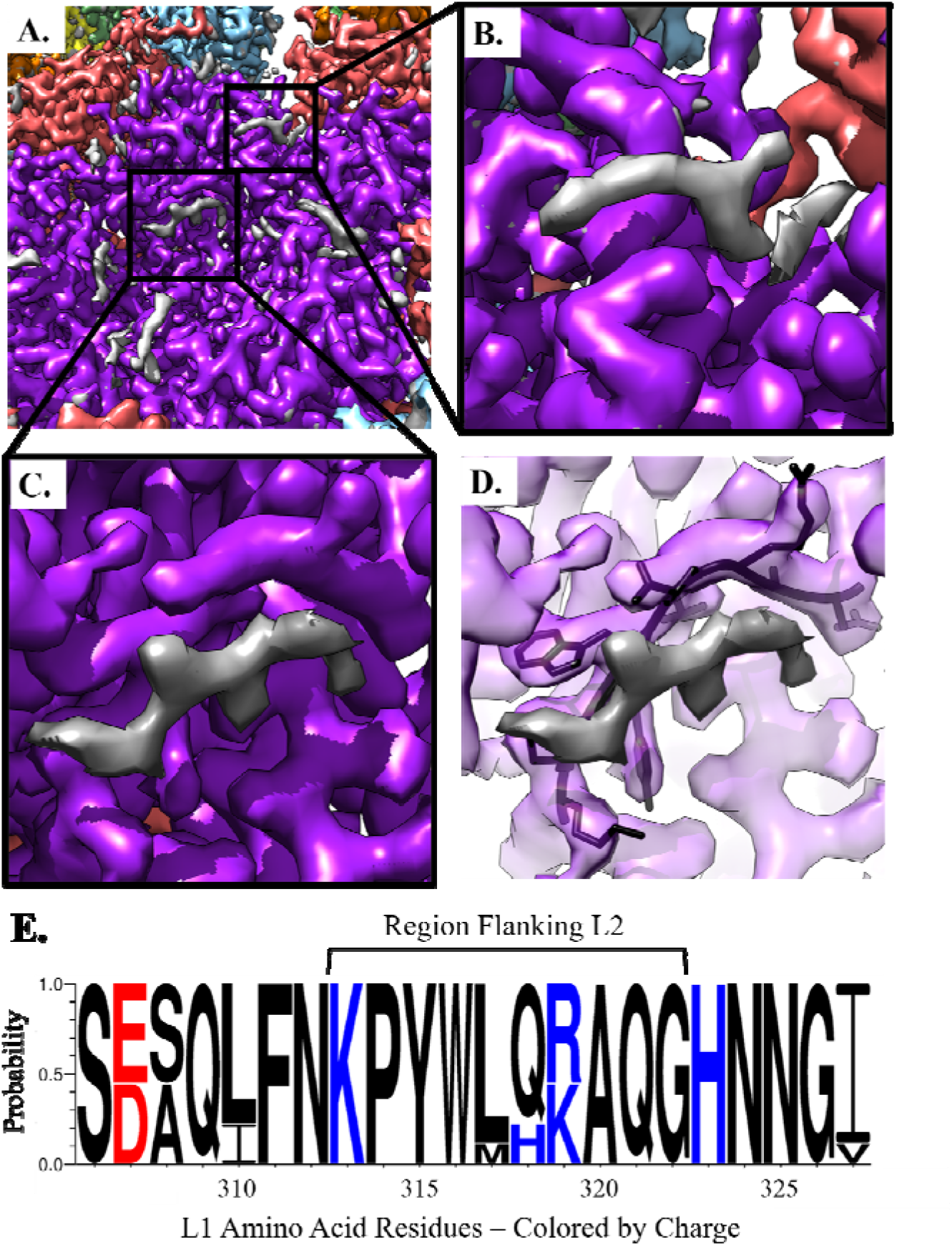
Non-L1 Internal Density. The capsomer density maps are colored within 2Å of the L1 protein chain and the gray areas are L2 density. (A) The internal capsomer core of pentavalent capsomer at contour level 2. The gray density is appears as protein-like density with two distinct areas correlated to each chain of L1 in the capsomer. (B) Zoom in of the fishhook L2 density. (C) Zoom in of internal capsomer density. (D) Same view as C. with L1 transparent, the residues that are being shown make up the conserved 306-loop, that is seen flanking the area of L2 density. (E) The plot shows the protein sequence conservation of the loop region Ser306 – Ile328. Colored by charge with red being acidic, and blue being basic. The internally exposed region is denoted by a bracket and is found on the interior of the capsomer and adjacent to non-L1 associated density.

The fragmented nature of the L2 density suggested the possibility of a symmetry mismatch between L2 and each L1 capsomer. Thus, to resolve L2 we attempted 3D classification of capsomers with and without symmetry expansion. However, no classes with distinct, continuous L2 density arose. This finding suggested two possibilities: (1) there was inadequate SNR for successful classification (2) L2 may have long disordered stretches between the locations that we were able to resolve.

### ISECC allowed correlated analysis of capsomers after local refinement

The original location of each extracted subvolume was recorded using ISECC (S. Table 3). Hexavalent capsomers were assigned a location relative to the nearest fivefold, threefold, and twofold symmetry axes, as well as a relative rotational assignment to distinguish the multiple capsomers most proximal to a given symmetry axis (S.Fig 4). These custom metadata were stored within the RELION 3.1 star file and after subvolume refinement the translational adjustments of each capsomer were correlated with respect to the other 71 capsomers, and to the capsid as a whole.

### Capsid diameter differences showed Gaussian distribution

The diameter of individual HPV particles was calculated using the capsomer metadata recorded by ISECC. Particle diameter was defined as the x,y distance along the icosahedral fivefold axis between pentavalent capsomers on opposite sides of the capsid. In order to minimize unmeasurable distances out of the image plane, this analysis was necessarily limited to particles where the polar-opposite fivefold capsomers lay within the same z plane, allowing a 5% radial tolerance (12.9Å) from z=0 (Fig 5.A). A diameter difference ratio was calculated for each qualifying capsid (Methods). Based on the icosahedrally averaged map diameter of 575Å, the 95^th^ percentile (0.988 – 1.017) corresponded to capsid diameters ranging from approximately 568 – 585Å, consistent with the variability of individual capsids observed in the micrographs (Fig 5.B, S.Fig 1.A). These results are more precise, but consistent with previous efforts to classify HPV into classes with discrete diameters(Guan et al., 2017b).

**Figure 5:**
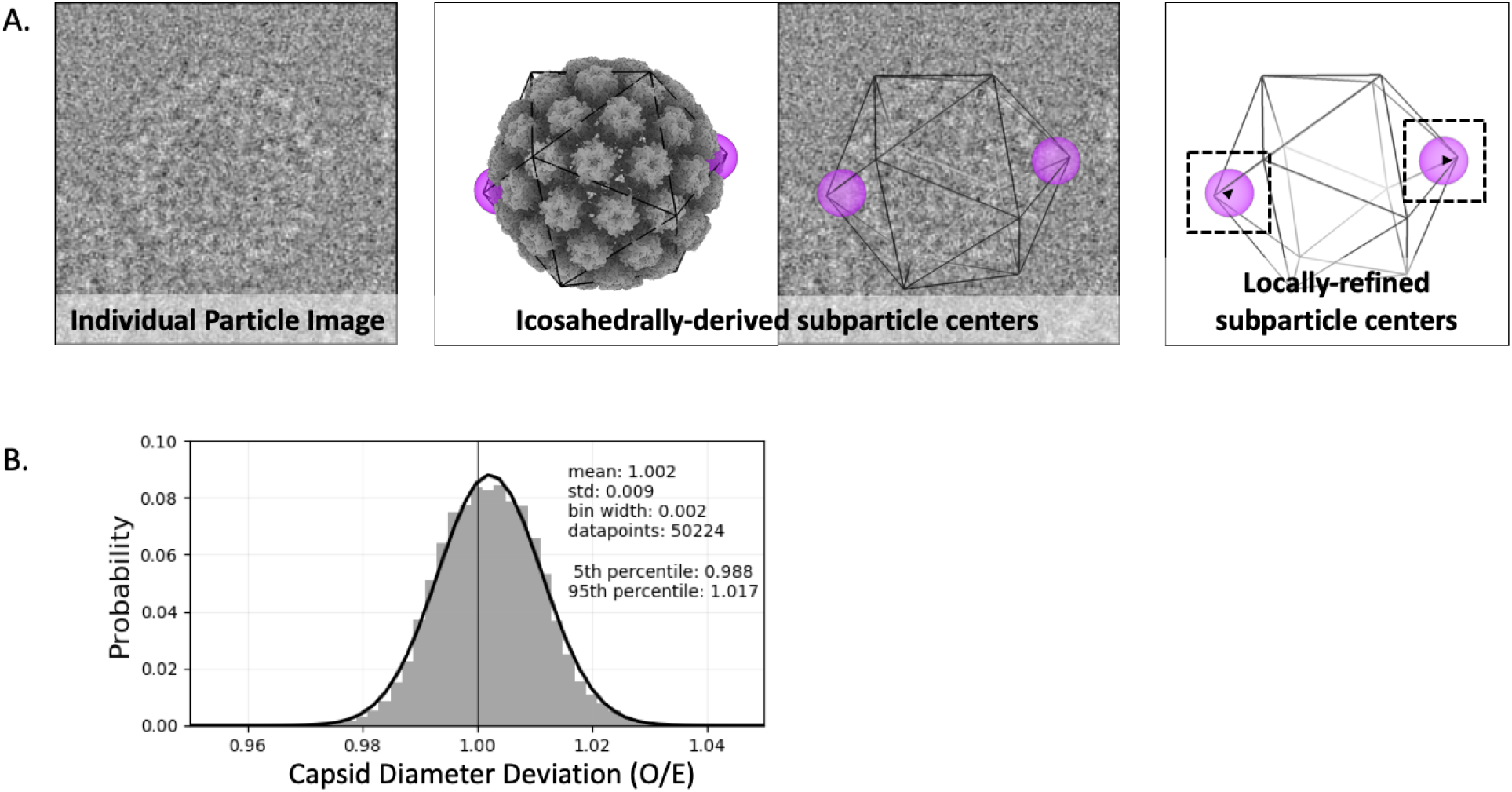
Capsid diameters. (A, left) From a cryo EM micrograph, an example of a particle oriented with opposing pentavalent capsomers in the Z=0 plane (within a tolerance, see Methods). (A, center) The same particle also illustrated in a 3D surface rendered model (gray) and outlined with icosahedral cage (black line) with opposing pentavalent capsomers indicated (purple spheres). The particle origin and orientation found during icosahedral refinement, determines the idealized capsomer locations. (A, right) Local subparticle refinement identifies the actual capsomer centers, which allows calculation of the deviation (black arrowheads) of the diameter along the given axis from expected values. (B) A gaussian distribution of capsid diameters (black curve) is evident after pooling all qualifying particles. A standard deviation of 0.9% corresponds to ~4.6Å, or 4.2 pixels at a pixel size of 1.1Å.

### Capsids have imperfect icosahedral symmetry

In addition to the improved resolution, local subvolume refinement allowed a determination of capsomer centers and orientations for the first time. The movement of capsomers relative to their icosahedrally-forced (idealized) parameters suggests flexibility. The corrected capsomer centers were used to evaluate the relative motions of capsomers within the imperfectly icosahedral HPV capsids. Capsomer movement was evaluated for all hexavalent subparticles within 25Å of x,y distance from the particle center. This criterion was selected to maximize lateral, in-plane movement of capsomers and limit out-of-plane motion, reflected in defocus, that cannot be recovered using local refinement. An additional filter was used to exclude any capsomer whose icosahedrally-derived center was within 15Å x,y distance of another, such as capsomers on the front and back of the capsid, as the identities of such capsomers could be confused during local refinement (Methods). For each qualifying hexavalent subparticle, the distances to all six neighboring capsomer centers were calculated based on their locally refined coordinates (Fig 6.A). These x,y distances were compared to the z-flattened icosahedrally averaged distances (Fig 6.B). The locally refined distances varied from the idealized values with a standard deviation of approximately 4%. Each of the four unique patterns of arm-exchange were defined for analysis as the hexavalent-pentavalent axis (A:F exchange); the fivefold-adjacent hexavalent-hexavalent axis (unidirectional B arm contribution); the threefold-adjacent hexavalent-hexavalent axis (C:D exchange); and the twofold hexavalent-hexavalent axis (E:E exchange). Each axis showed a similar pattern of deviation with expansion or contraction of 6-7% (95^th^ percentile range). There was no obvious correlation of distance deviations for a given grouping of capsomers (e.g. all contracted or all expanded).

**Figure 6:**
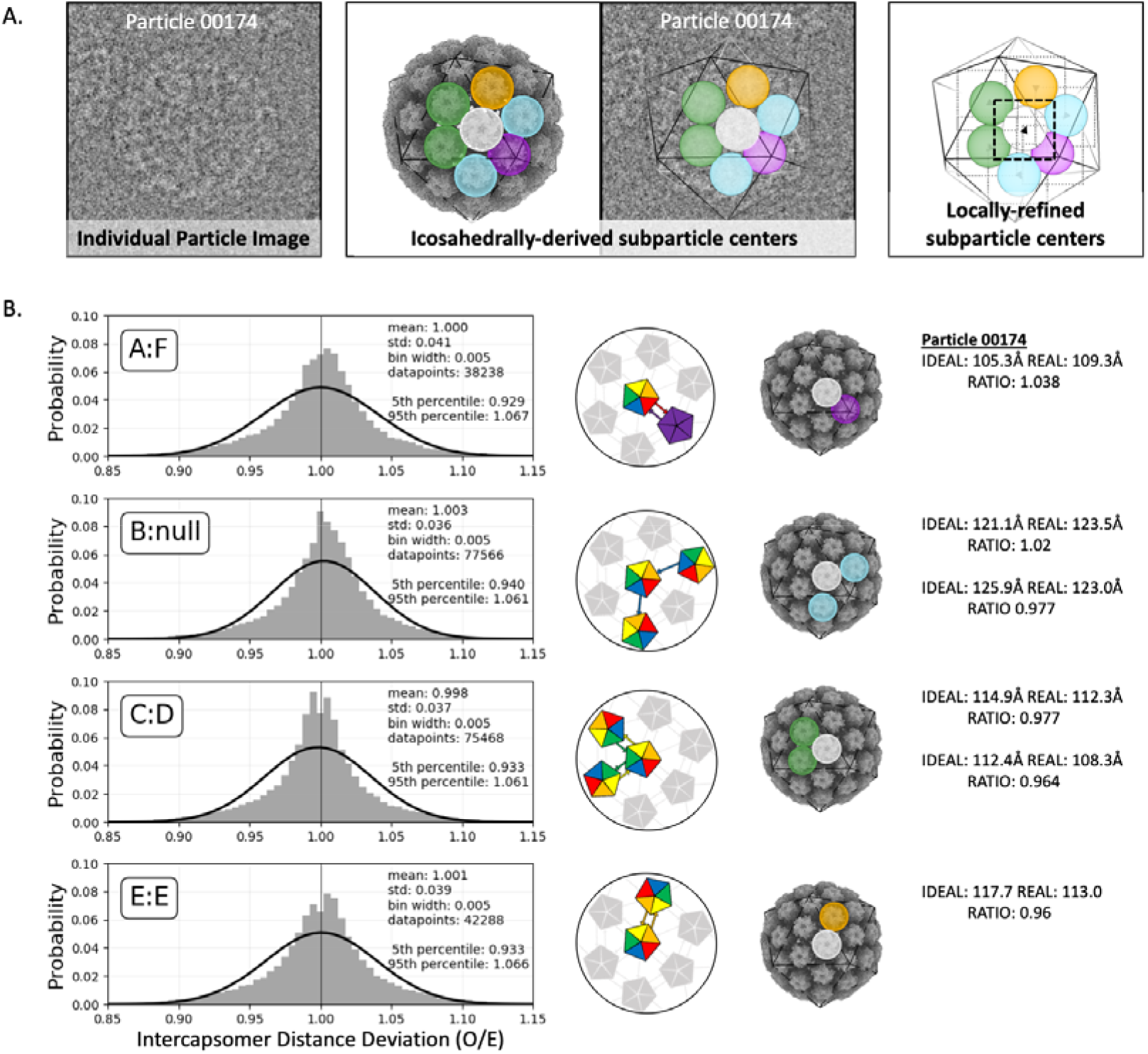
Inter-capsomer distances. (A, left) An example of a particle oriented in one of the cryo EM micrographs so that it met the selection criteria (Methods) for capsomer distance analysis. (A, center) The same particle also illustrated in a 3D surface rendered model (gray) and outlined with icosahedral cage (black line) with a reference capsomer (white sphere) indicated centrally in the hexavalent capsomer. Relative to the reference, there are four different arm configurations: the hexavalent-pentavalent axis (A:F arm exchange) (purple); the fivefold-adjacent hexavalent-hexavalent axis (unidirectional B arm contribution) (blue); the threefold-adjacent hexavalent-hexavalent axis (C:D exchange)(green); and the twofold hexavalent-hexavalent axis (E:E exchange)(orange). The particle origin and orientation found during icosahedral refinement determined the idealized capsomer locations. (A, right) Local subparticle refinement found the correct capsomer centers (black arrowhead), which then can be used to determine the deviation from the icosahedrally-derived, idealized parameters. (B) Deviation of inter-capsomer distances from idealized values were plotted and the ratios corresponded to contraction (values < 1) or expansion (> 1). The four unique axes of arm configurations as defined above were plotted. Each axis shows a similar, non-gaussian pattern of deviation (gaussian fit, black curve) with a standard deviation of ~4%. Values and schematic for the example particle are shown at right.

Flexibility analysis suggested that variation in capsid diameter is due to variable contraction or expansion of the flexible C-terminal arms linking the capsomers, which in turn move as rigid bodies. This is consistent with the poor resolution of residues 431-439, found in the connecting arms, that was seen in all chains of the asymmetric unit in both the icosahedrally refined and locally refined maps. However, we saw no evidence suggesting that any one capsomer-to-capsomer linkage is more or less flexible than another. This observation is surprising considering some capsomers are paired by donation of a single arm (chain B:Null), whereas others are linked by bidirectional arm exchange (chain E:E, A:F, & C:D) (Fig 6.B).

## Discussion

Previous icosahedral reconstructions of papillomaviruses never surpassed ~4.3Å resolution(Guan et al., 2017b). This resolution limitation cannot be directly attributed to the use of VLPs with unauthentic genome (quasi or pseudovirus), as studies with native papillomavirus have encountered the same resolution barrier(Wolf et al., 2010). Here we show that even with recent improvements in cryo EM hardware and software for whole particle sorting, classification, and averaging, the resolution has not improved. However, we found that HPV responded well to a subparticle refinement approach that isolated individual capsomers for local refinement. Thus, we were able to improve the resolution from 4.5Å to 3.1Å. The maps of individual capsomers were subsequently recombined to create a 3.1Å resolution model of the complete icosahedral virus capsid.

The high resolution and the structure of the L1 asymmetric unit allowed us to better understand the different conformations of the connecting arms that link capsomers. Although each L1 chain is chemically identical, each 37 amino acid loop makes a different connection to its neighboring capsomer, due to quasi-equivalence. When looking at the six chains of the asymmetric unit, the low RMSD of the L1 capsomer cores, compared to the higher RMSD of the connecting arms, suggests that the L1 core secondary structure is conserved whereas the arms have different conformations. One thing that is consistent in all chains of L1 within the asymmetric unit is the modest resolution of His431 to Asp439, which are solvent exposed on the surface of the virus, and the poor resolution is likely attributed to those nine amino acids being a source of flexibility.

The near atomic resolution of the map allowed us to identify protein-like density unfilled by L1, which corresponded to the minor capsid protein, L2. In our map we can see clear separation between the L1 chain and regions of L2, which have likely been stabilized by their interactions with L1. Notably, there is a high sequence conservation of the flanking L1 residues that interact with L2 in these regions. However, the L2 density is not as strong as L1 in either location although the magnitude of L2 density is stronger in the pentavalent than the hexavalent capsomer (S.Fig 3). Size constraints of the capsomer pore would prevent incorporation of five copies of L2 in any one capsomer thus indicating a symmetry mismatch between L1 and L2. We estimate that one or two L2 proteins could fit under each capsomer, consistent with the L2 density magnitude compared to L1. Unfortunately, multiple attempts at 3D classifications failed to solve a symmetry mismatch between L1 and L2 and produce continuous density between the L2 fragments that we see. This implies L2 may possess significant disordered stretches between the resolved regions that are ordered by L1 interactions.

Variability in previous L2 stoichiometry studies suggests that L2 is probably asymmetrically incorporated within the capsid and there may be a range in copy number of L2 proteins per individual capsid within a population. Our results suggest that L2 can be incorporated into the core of any capsomer, but is more prevalent under the pentavalent capsomers, as indicated by stronger density in these capsomer environments. However, we cannot rule out that L2 is missing altogether from some capsomers. The wide variability prevents an estimation of the stoichiometry of L2 which may differ among capsids as discussed above. Our ability to resolve only small segments of L2 further suggests that the overall structure of L2 is largely disordered, with resolved segments ordered by their interactions with L1.

Different HPV capsid diameters are obvious by eye in the micrographs and have been recorded previously by Guan et al. 2017, but attempts to overcome the heterogeneity by sorting into discrete classes, failed to achieve better than 4.3Å resolution(Guan et al., 2017b). Previously, heterogeneity was attributed to the maturation of the capsid, specifically in the disulfide bonds of the C-terminal extensions (connecting arms) between capsomers(Guan et al., 2017b). Because subparticle refinement was able to determine the true center of each capsomer, we were able to use these new values to determine the actual diameter of each capsid on a per particle basis using ISECC_local_motions.py script (Methods). Here, we demonstrated a measurable contraction and expansion of capsids relative to the icosahedrally-enforced average. The 95^th^ percentile encompasses a range from −1.2 to 1.7% (17Å), covering 15.5 pixels under our imaging conditions. This range is broad enough that the observed variation cannot be attributed solely to optical aberrations, imprecise Euler angle assignment, or reference-drift during subparticle refinement. Notably, refinement of higher order aberrations, magnification anisotropy, and per-particle defocus parameters failed to improve resolution during whole particle icosahedral refinement.

Additionally, the metadata labels implemented in ISECC were used to track deviations in the location of discrete capsomers with respect to their neighbors compared to their expected location in perfect icosahedral symmetry. We showed that inter-capsomer distances, governed by the connecting arms, can expand or contract by up to 7% (95^th^ percentile, range of about 7-9Å in either direction) from their idealized values, without bias in the magnitude or direction of capsomer movement (Fig 6). This flexion between capsomers provides the basis for the capsid-wide variability seen in the diameter analysis.

Although dynamics information cannot be conclusively derived from a vitrified population, each particle may represent a state within an equilibrium of particles in solution. These states, described for the first time by ISECC, provide a source of experimental capsid dynamics data that could contribute to whole-capsid molecular dynamics simulations such as described by Hadden *et al.* in hepatitis B virus(Hadden et al., 2018). We propose a model in which the HPV capsid is dynamic and in a state of constant flexing. This model is supported by the composition of stable beta-jellyroll capsomer cores with flexibility stemming from the connections between these capsomers. The classification of different diameter particles likely failed because the intercapsomer flexing is global and variable, not by symmetrical coordinated movements. Thus, even for particles with the same diameter, the capsids have different configurations due to the flexing between capsomers that creates imperfect icosahedral symmetry. These elements of heterogeneity include a continuous range of different diameters due to deviations of capsomer positions from a perfect icosahedral grid, which has likely limited the resolution attained through whole-capsid refinement.

The observed dynamics may be a natural consequence of the architecture of the T=7d icosahedral capsid that requires L1 to exist in hexavalent and pentavalent environments. However, there are a multitude of possible benefits to a dynamic capsid that allows reversible expansion and contraction. This flexing could serve as a defense mechanism to hide or alter antibody binding sites, inhibiting recognition and preventing binding and neutralization. Capsid flexibility also has the potential to make HPV more resilient to environmental factors such as pH, temperature, or desiccation. Mechanistically, dynamics are certainly necessary for the conformational changes that are known to occur during host cell entry. Lastly, in the final stages of the virus life cycle, mature virions pack within the nucleus in a paracrystalline array of compact capsids that are significantly smaller diameter than when observed after lysis in suspension. Thus, capsid flexibility is an intriguing and essential factor to papillomavirus function.

Until now, high resolution structures of HPV have been unattainable. By using subparticle refinement approaches we demonstrated that high resolution is achievable. The new, more accurate L1 structure revealed L2 density flanking the conserved 306-loop of L1. We also found that the HPV capsids are globally flexible with capsomers moving as rigid bodies. These findings provide a framework for continuing structural, biochemical, genetic, and biophysical studies of HPV. These new observations on capsid flexibilities containing rigid capsomer units paves the way for design of HPV capsid platforms that can deliver foreign antigenic epitopes engineered for the development of second-stage vaccines (e.g. L2 insertions into L1 sequences, Schellenbacher *et al.*, 2009)(Schellenbacher et al., 2009). The design of stable antigenic epitope-expressing HPV particles will be improved with the cryo EM structural analyses presented in this study, and lead to *in silico* rational design and selection of stable VLP vectors for vaccines and gene-therapy deliverables.

## Materials & Methods

### Preparation of Virus

HPV16 quasivirus containing L1 and L2 proteins and encapsidating a CRPV genome having the SV40 origin of replication was prepared as described previously(Brendle et al., 2010; Mejia et al., 2006; Pyeon et al., 2005). In brief, HPV16 sheLL plasmid (kindly provided by John Schiller, NIH) was transfected together with linear CRPV/SV40ori DNA into 293TT cells and prepared as described previously(Buck, Pastrana, et al., 2005; Buck, Thompson, et al., 2005; Pastrana et al., 2004). HPV16 was allowed to mature and then pelleted by centrifugation. The centrifuged pellet was resuspended in 1 M NaCl and 0.2 M Tris (pH 7.4). After CsCl gradient purification, the lower band was collected. The lower band was then added to CsCl for another round for ultracentrifugation to separate full from empty virus particles. The lower band was collected, concentrated, and buffer exchanged using a 100KDa cutoff spin column as described previously(Guan, Bywaters, Brendle, Lee, Ashley, Makhov, et al., 2015; Guan et al., 2017b). The concentrated HPV16 quasivirus particles were applied to 300 mesh carbon coated copper grids and stained with 2% phosphotungstic acid. The sample was analyzed for integrity and concentration on an FEI Tecnai G2 Spirit BioTwin transmission electron microscope.

### Cryo-EM Data Collection

The HPV16 sample was assessed for purity and concentration before vitrification for cryo-EM data collection on the Penn State Titan Krios (https://www.huck.psu.edu/core-facilities/cryo-electron-microscopy-facility/instrumentation/fei-titan-krios). 3.5 microliters of the purified virus sample was pipetted onto a glow-discharged R2/1 Quantifoil grid (Quantifoil Micro Tools GmbH, Jena, Germany), blotted for 2.5 seconds, and plunge-froze in liquid ethane using a Vitrobot Mark IV (Thermo Fisher, USA). Vitrified grids were imaged with the use of a Titan Krios G3 (Thermo Fisher, USA) under automated control of the FEI EPU software. An atlas image was taken at 165x magnification, and suitable areas were selected for imaging on the FEI Falcon 3EC direct electron detector. The microscope was operated at 300 kV with a 70 μm condenser aperture and a 100 μm objective aperture. Magnification was set at 59,000x yielding a calibrated pixel size of 1.1 Å. Four, nonoverlapping exposers were acquired per each 2-um-diameter hole of the grid with the beam in parallel mode, for an overall collection of 10,143 micrographs. The total dose per exposure was set to 60 e^−^/Å^2^ (Supp. Table 1).

### Icosahedral Refinement

Icosahedral refinement was performed in cryoSPARC(Punjani et al., 2017). The micrographs underwent full frame motion correction and CTF estimation (CTFFIND4)(Rohou & Grigorieff, 2015). Micrographs were curated and sorted to reject micrographs with crystalline ice. Particles were picked using 2D templates from 840 particles. Local motion correction was performed on the particle stack and the CTF estimated micrographs. The particles went into a homogenous refinement. The final resolution was determined by gold standard FSC threshold of 0.143.

### Icosahedral Subparticle Extraction and Correlated Classification

ISECC_subparticle_extract was used after icosahedral refinement to divide each particle image into subparticles. Both hexavalent and pentavalent capsomer subparticle were separately created. An initial model was made for each subparticle type using relion_reconstruct with 10,000 subparticles(Scheres, 2012). The subparticles were then processed in RELION v3.1(Scheres, 2012).

### Local Subparticle Refinement

Pentavalent subparticles were locally refined with C5 symmetry whereas hexavalent C1 was used to refine the pentavalent subparticles. Each subparticle dataset was refined with a spherical mask of 200Å applied to focus on the capsomer and adjoining arms. The final resolutions were determined by gold standard FSC with a threshold of 0.143 in RELION post-process. Local resolution maps were generated using RELIONs own software.

### Icosahedral recombined map

Postprocessed subparticle maps were recombined into a complete capsid using ISECC_recombine. This procedure is similar to the recombination process for subparticles in Block Based Reconstruction and LocalRec (Abrishami et al., 2020; Ilca et al., 2015; Zhu et al., 2018).

Briefly, ISECC_recombine loads the subparticle maps into a numpy array and both shifts and rotates the maps to their locations in an idealized icosahedron, using regular grid interpolation in real space. This interpolation scheme allows merging of the subparticle-refined models into a single asymmetric unit.

### Model Building

The previously solved L1 structure (PDB: 5KEP) was used to initiate the build(Guan et al., 2017b).

The model was visually inspected and adjusted during iterative refinement before a final validation(Adams et al., 2010; Emsley et al., 2010). The L1 protein structure from HPV16 (PDB: 5KEP) was used as an initial model. The asymmetric unit PDB was used to create a protein structure for the pentavalent and hexavalent capsomers. The hexavalent capsomer was made from chains B-F of the existing PDB and the pentavalent capsomer model was made from 5 copies of chain A from the asymmetric unit. These models were fit into the electron density map of the pentavalent and hexavalent capsomers independently in Chimera(Pettersen et al., 2004). The protein structure of L1 was then refined in real space against the cryo-EM electron density map in Phenix with geometry and secondary structural restraints(Adams et al., 2010). The structure was visually inspected and manually refined in Coot and validated using MolProbity(V. B. Chen et al., 2010; Emsley et al., 2010). (Supp. Table 1) The protein structures RMSD values were calculated using MatchMaker in Chimera(Pettersen et al., 2004).

### Icosahedral Subparticle Extraction & Correlated Classification (ISECC)

ISECC is a Python-based subparticle extraction package inspired by localized reconstruction and block-based reconstruction, compatible with RELION 3.1(Ilca et al., 2015), (Zhu et al., 2018). This allows the user to take advantage of higher order aberration correction within the updated RELION CtfRefine pipeline.

Subparticles are generated according to the following parameters: --vector (atomic coordinate, in angstroms, for the subparticle center), --roi (region of interest: fivefold, threefold, twofold, or fullexpand), --subpart_box (box size for the subparticle images, in pixels), -- supersym (symmetry of the whole particle: I1 or I2).

ISECC introduces several new metadata labels to the RELION 3.1 star file to enable correlated analysis of subparticles after local refinement or classification. For the idealized icosahedron, each vertex is given a designation, rlnCustomVertexGroup, containing the symmetry axis and an integer (e.g., 5f01) (SFig 4). If the chosen subparticle is off a strict symmetry axis, rlnCustomVertexGroup is defined according to the nearest three symmetry axes, e.g., 5f07.3f14.2f20. Given that each symmetry axis has multiple associated subparticles (5 per fivefold, etc.), each subparticle is additionally given a rotational specifier (a-e, a-c, or a-b), ordered anti-clockwise about the local symmetry axis. As such, the complete rlnCustomVertexGroup specifier for an off-axis subparticle takes a form such as 5f07e.3f14c.2f20a. This allows grouped analysis of, for example, all five subparticles belonging to local vertex 5f07 for a given viral particle.

Metadata label rlnCustomOriginXYZAngstWrtParticleCenter contains information on the icosahedrally refined offset of each subparticle in X,Y,Z with respect to the whole particle center. rlnCustomRelativePose contains the relative pose of the subparticle with respect to the icosahedrally refined orientation. rlnCustomRelativePose is stored in quaternion format rather than the standard RELION Euler angles, rlnAngleRot, rlnAngleTilt, rlnAnglePsi.

### Local Refinement of Pentavalent and Hexavalent Capsomers

Pentavalent and hexavalent capsomers were extracted in ISECC using the following commands, respectively:

> [--vector 0 136 220 --roi fivefold --supersym I1 --subpart_box 300]

and,

> [--roi fullexpand --supersym I1 --subpart_box 300 --vector 42 42 259 --batchsize 9000]

Capsomers were locally refined in RELION 3.1 using the following commands:

> [‘which relion_refine_mpi --o Refine3D/job036/run --auto_refine --split_random_halves --i fivefold_subparticles/fivefold_20200114_1026/fivefold_subpart _PRIOR.star --ref fivefold_subparticles/fivefold_20200114_1026/fivefold_initialmodel_c5.mrc -- firstiter_cc --ini_high 20 --dont_combine_weights_via_disc --scratch_dir/scratch/sxh739/ --pool 100 --pad 2 --ctf --ctf_corrected_ref --particle_diameter 200 -- flatten_solvent --zero_mask --oversampling 1 --healpix_order 5 -- auto_local_healpix_order 5 --offset_range 3 --offset_step 2 --sym C5 -- low_resol_join_halves 40 --norm --scale --j 1 --gpu “” --dont_check_norm --sigma_ang 1.5 --pipeline_control Refine3D/job036/]

and,

> [‘which relion_refine_mpi’ --o Refine3D/job040/run --auto_refine --split_random_halves --i fullexpand_subparticles/fullexpand_20200123_1329/fullexpand_subpart_PRIOR.star --ref fullexpand_subparticles/fullexpand_20200123_1329/fullexpand_initialmodel_c1.mrc -- firstiter_cc --ini_high 20 --dont_combine_weights_via_disc --scratch_dir/scratch/sxh739/ --pool 100 --pad 2 --ctf --ctf_corrected_ref --particle_diameter 200 -- flatten_solvent --zero_mask --oversampling 1 --healpix_order 5 -- auto_local_healpix_order 5 --offset_range 3 --offset_step 2 --sym C1 -- low_resol_join_halves 40 --norm --scale --j 1 --gpu “” --dont_check_norm --sigma_ang 1.5 --pipeline_control Refine3D/job040/]

### Correlation of locally refined capsomers coordinates

Correlational analysis was performed using the ISECC_local_motions script. Briefly, this parses the locally refined pentavalent and hexavalent capsomer star files to evaluate local deltas for subparticle origins and poses as compared to their idealized, icosahedrally derived, starting values. Coupled with the new metadata identifiers, rlnCustomVertexGroup and rlnCustomOriginXYZAngstWrtParticleCenter, this identifies deviation of each capsomer from idealized icosahedral symmetry on a per-particle basis.

Deviation in particle diameter 3was calculated for all particles that satisfied selection criteria, namely, a pair of pentavalent capsomers within +/− 5% (12.9Å) of the central plane (Z=0), where z_max_ corresponded to the particle radius as defined by the distance between the center of capsomer and the particle center. This geometry minimizes contribution to distance along the Z-axis, which unlike X or Y cannot be locally refined. These capsomers are easy identified using the Z parameter within rlnCustomOriginXYZAngstWrtParticleCenter. The locally refined XY distance between polar opposite pentavalent capsomers was calculated for the 50,224 qualifying particles and compared to the Z-flattened icosahedrally-derived distance, producing a difference ratio. It is important to note that this analysis cannot capture the Z-component of any diameter deviation, necessitating the selection criteria described above.

Capsomer distance analysis was conducted on all particles with a hexavalent subparticle center within 25Å x,y distance of the whole particle center. Capsomers with a potential doppelganger within 15Å x,y distance on the opposite face of the capsid were excluded from this analysis to avoid the risk of subparticle identity swap during local refinement. Neighboring subparticles were then identified using the rlnCustomVertexGroup parameter, allowing the geometric relationships between capsomers to be distinguished (see the four relationships listed in Fig 6b). The locally-refined x,y distance for each pair of qualifying capsomers (A:F 38,238, B:Null 77,566, C:D 75,468, E:E 42,228) was divided by the icosahedrally-derived x,y distance (dropping the z coordinate) to produce an in-plane ratio corresponding to contraction or expansion along the given axis.

## Acknowledgements

This work was supported in part by the J. Gittlen Memorial Golf Tournament and the Pennsylvania Department of Health CURE funds.

## Author Contributions

DJG, SRH, NDC, and SLH conceived the study. SS expressed and purified virus. CMB vitrified the sample and collected cryo EM data. DJG designed and developed the custom software. SRH solved the structures and built the models. DJG, SRH, and SLH interpreted the data. DJG, SRH, NDC, and SLH wrote the manuscript.

## Data and Code Availability

The HPV structures of the 3.1Å recombined map (EMDB: XXXX), pentavalent capsomer map (EMDB: XXXX), hexavalent capsomer map (EMDB: XXXX), and 4.4Å icosahedral refinement (EMDB: XXXX) have been deposited in the EM database (http://www.emdatabank.org/). Coordinates for the atomic model of the asymmetric unit of HPV16 L1 proteins (PDB: XXXX) have been deposited in the protein data bank (https://www.rcsb.org/). ISECC, our custom software for subparticle extraction and correlated classification, is available on GitHub (https://github.com/goetschius/isecc)

## Supplementary Information

**SFig 1.**
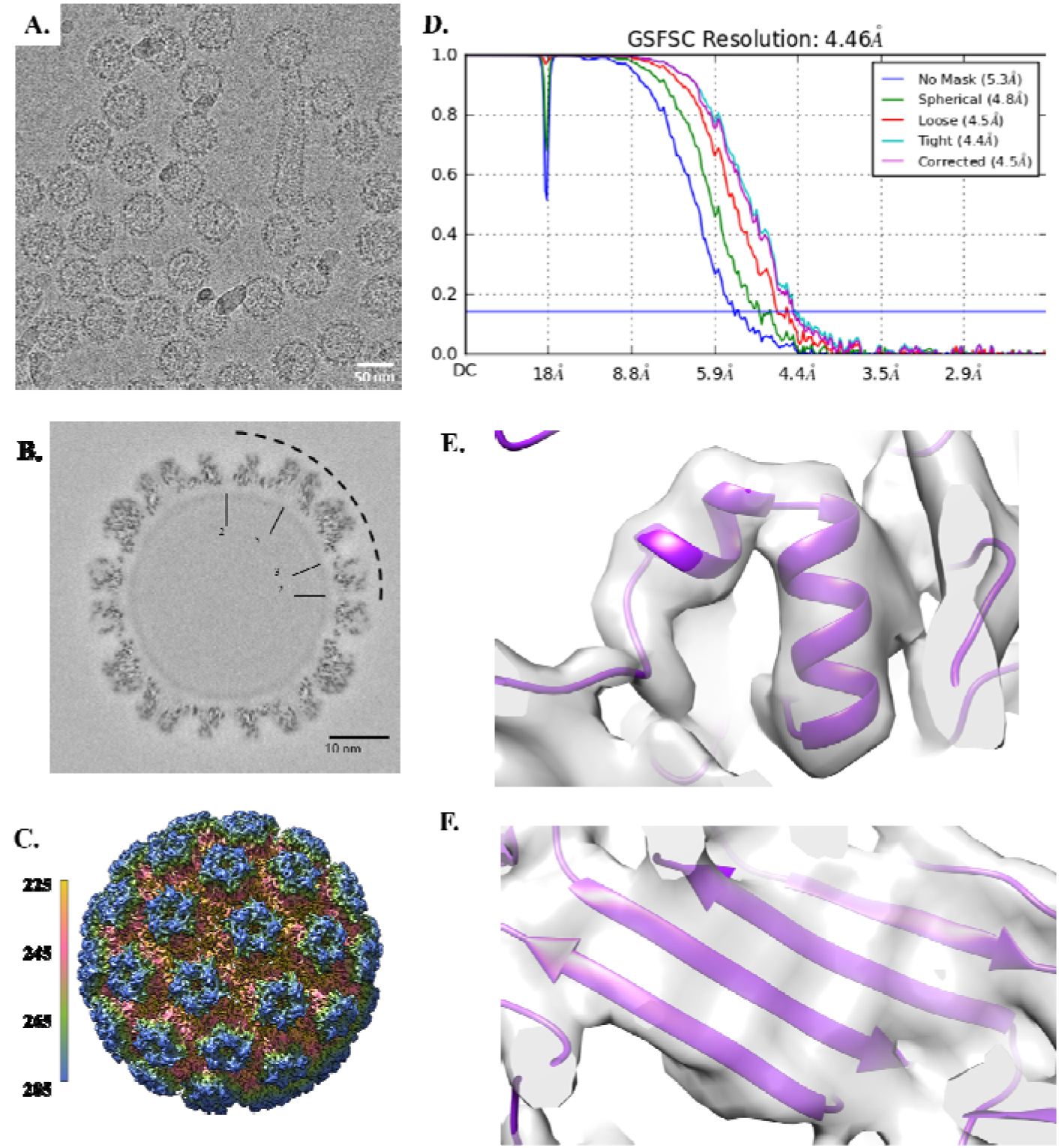
Icosahedral Reconstruction. (A) Representative micrograph 1,164 out of 8,936 collected. (B) Central cross-section of icosahedral structure with the symmetry labeled in the dotted region. (C) Full icosahedral reconstruction colored radially in Angstroms. (D) FSC Curve for icosahedral refinement. (E) Representative alpha helix (amino acid residues: 385-394, 396-401) at 4.5Å. (F) Representative beta sheet (amino acid residues: 71-76, 335-325, 153-160, 254-248) at 4.5Å.

**SFig 2.**
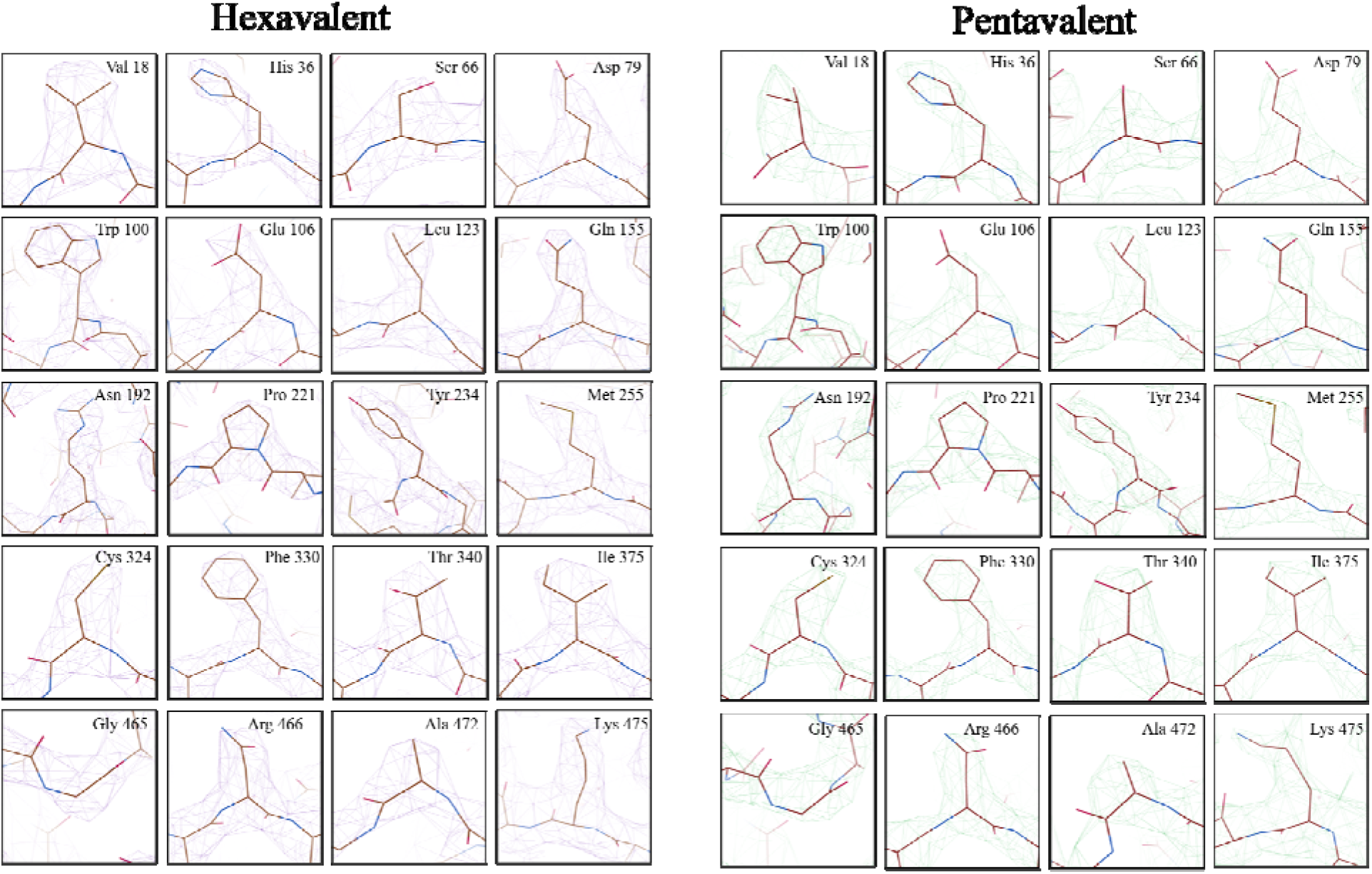
Representative density of all side chains in pentavalent and hexavalent capsomers. Representative side chains were chosen from L1 (brown) and compared between the pentavalent (purple density) and hexavalent (green density) capsomer environments.

**SFig 3.**
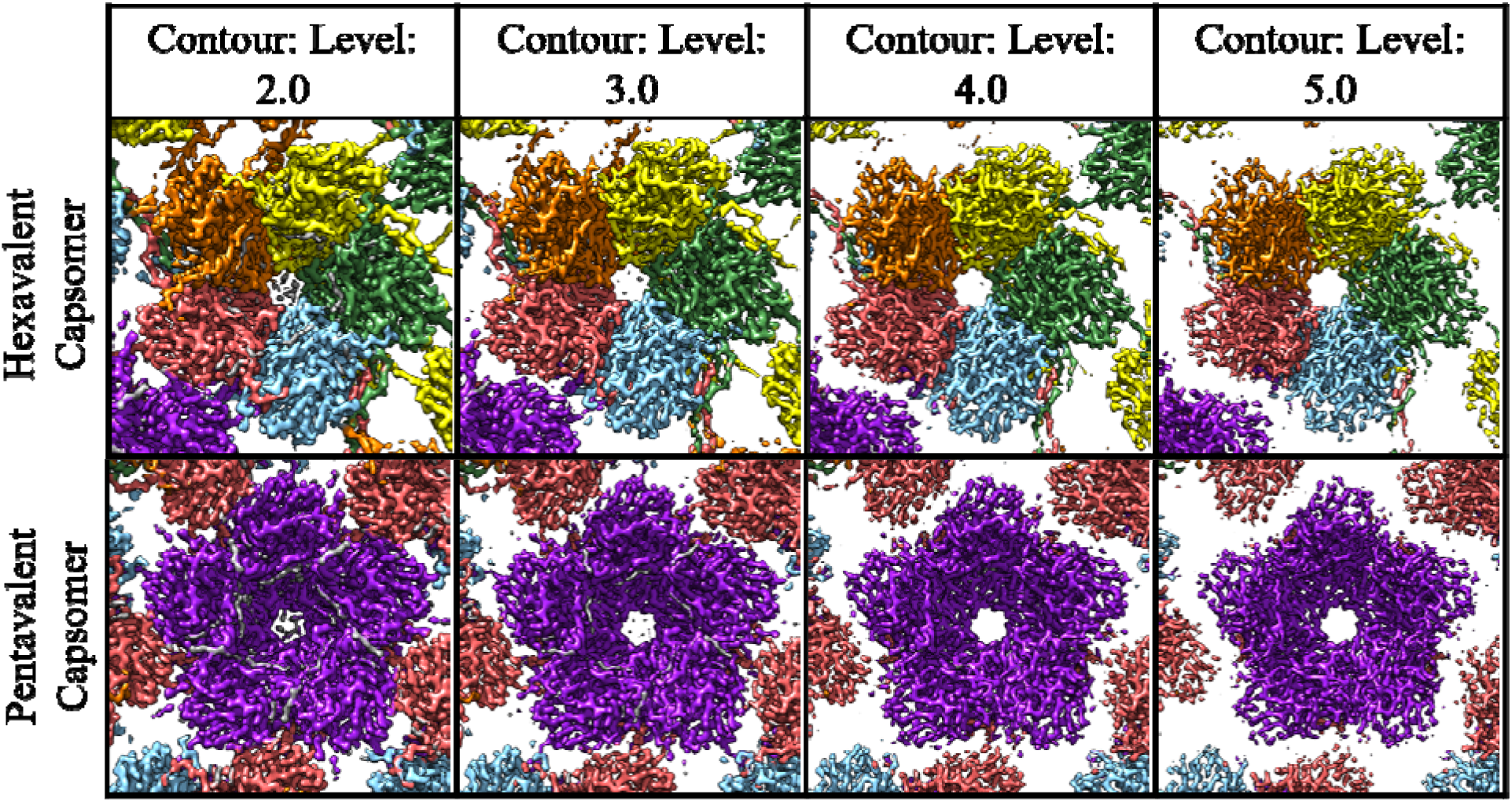
L2 Density. The surface rendered hexavalent (top) and pentavalent (bottom) capsomer density maps were colored as in Fig. 1A within 2Å of the L1 protein chain with unfilled density (gray) corresponding to L2. The L2 density is stronger in the pentavalent capsomer as can be seen with the changing contour of the map. In the hexavalent capsomers the same internal density can be noted, but the density disappears along with noise. Two distinct areas of gray density can be seen that are correlated to each chain of L1 in the capsomer.

**SFig 4.**
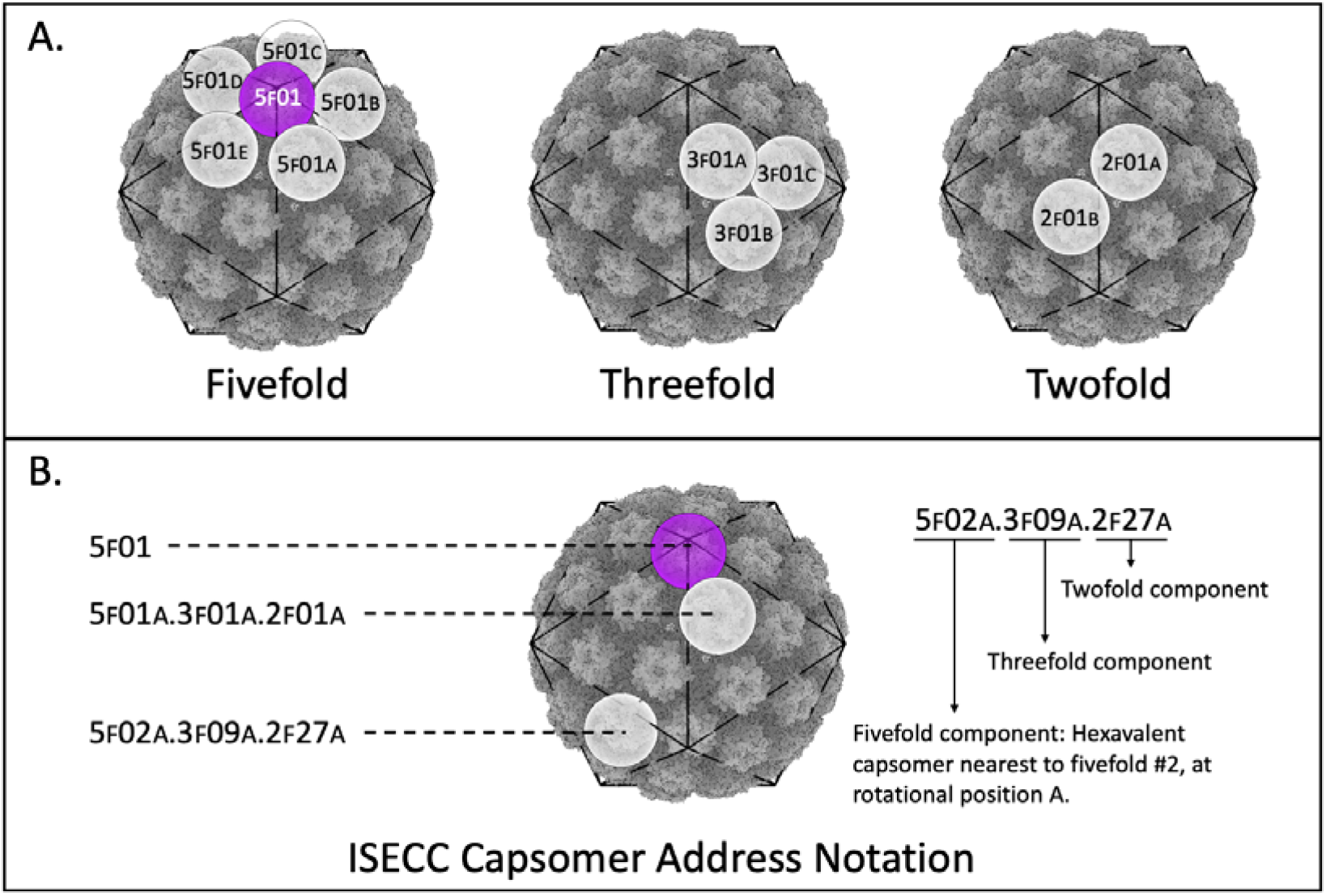
Capsomer addresses assigned in ISECC. A) Each capsomer is assigned an address identifying the nearest symmetry vertices. Pentavalent capsomers only have a fivefold designation without any rotational parameter (left, purple). Hexavalent capsomers receive designations for the nearest fivefold (left), threefold (center), and twofold (right) axis, as well as rotational parameters. B) Examples of complete addresses implemented in ISECC are shown as one-part (pentavalent) and three-part (hexavalent) capsomer designations. Addresses are assigned during subparticle generation after normalization of the input vectors to a standard, shared asymmetric unit. This allows refinement parameters for any given subparticle to be correlated with other subparticles from the same parental particle.

**Supp. Table 1.**
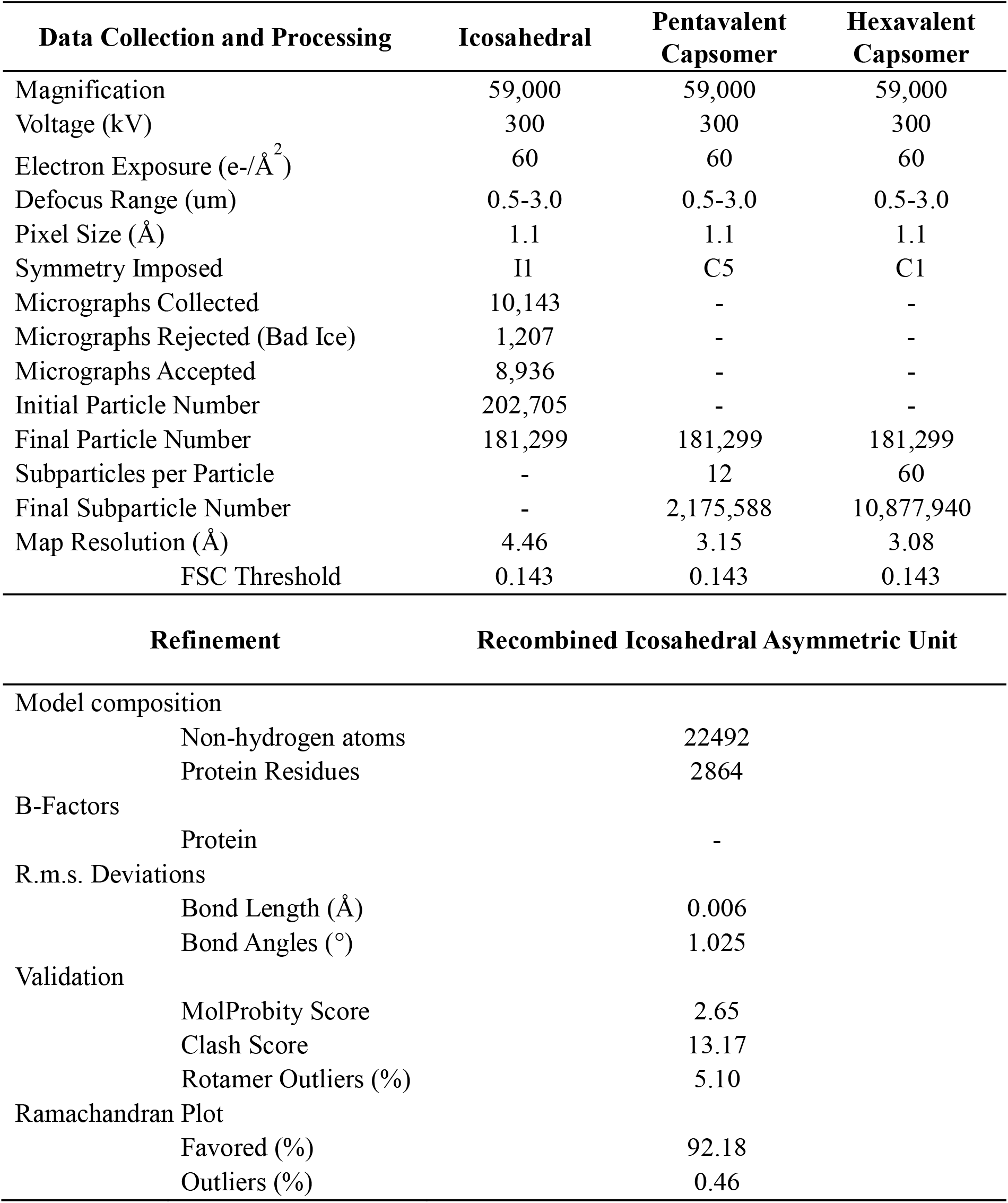
Cryo-EM data collection, refinement and validation statistics.

**Supplemental Table 2.**
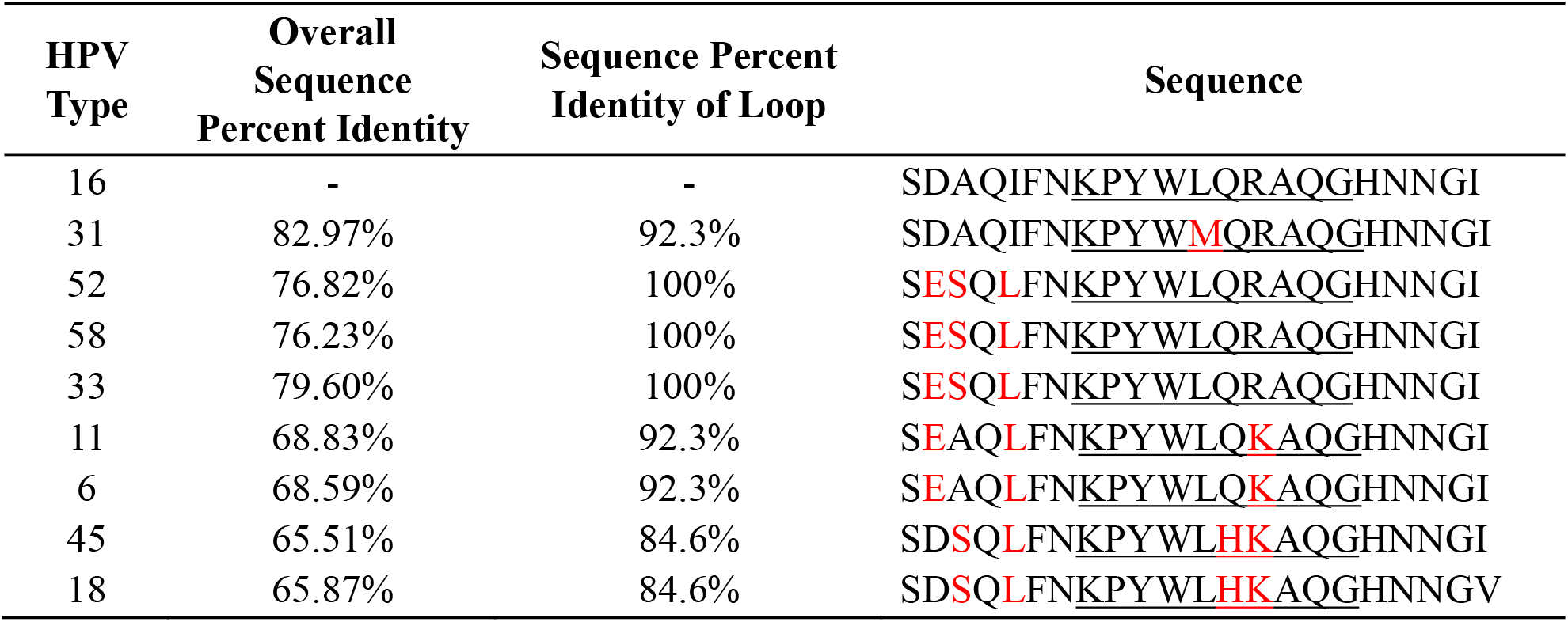
Sequence Alignment of Ser306 – Ile328 Loop Region.

| HPV Type | Overall Sequence Percent Identity | Sequence Percent Identity of Loop | Sequence |
| --- | --- | --- | --- |
| 16 | - | - | SDAQIFNKP<u>Y</u>WLQRAQGHNNGI |
| 31 | 82.97% | 92.3% | SDAQIFNKP<u>Y</u>W<u>M</u>QRAQGHNNGI |
| 52 | 76.82% | 100% | **SE**SQ**L**FNKP<u>Y</u>WLQRAQGHNNGI |
| 58 | 76.23% | 100% | **SE**SQ**L**FNKP<u>Y</u>WLQRAQGHNNGI |
| 33 | 79.60% | 100% | **SE**SQ**L**FNKP<u>Y</u>WLQRAQGHNNGI |
| 11 | 68.83% | 92.3% | **SE**AQ**L**FNKP<u>Y</u>WLQ**K**AQGHNNGI |
| 6 | 68.59% | 92.3% | **SE**AQ**L**FNKP<u>Y</u>WLQ**K**AQGHNNGI |
| 45 | 65.51% | 84.6% | SD**S**Q**L**FNKP<u>Y</u>WL**H**KAQGHNNGI |
| 18 | 65.87% | 84.6% | SD**S**Q**L**FNKP<u>Y</u>WL**H**KAQGHNNGV |

**Supplemental Table 3:**
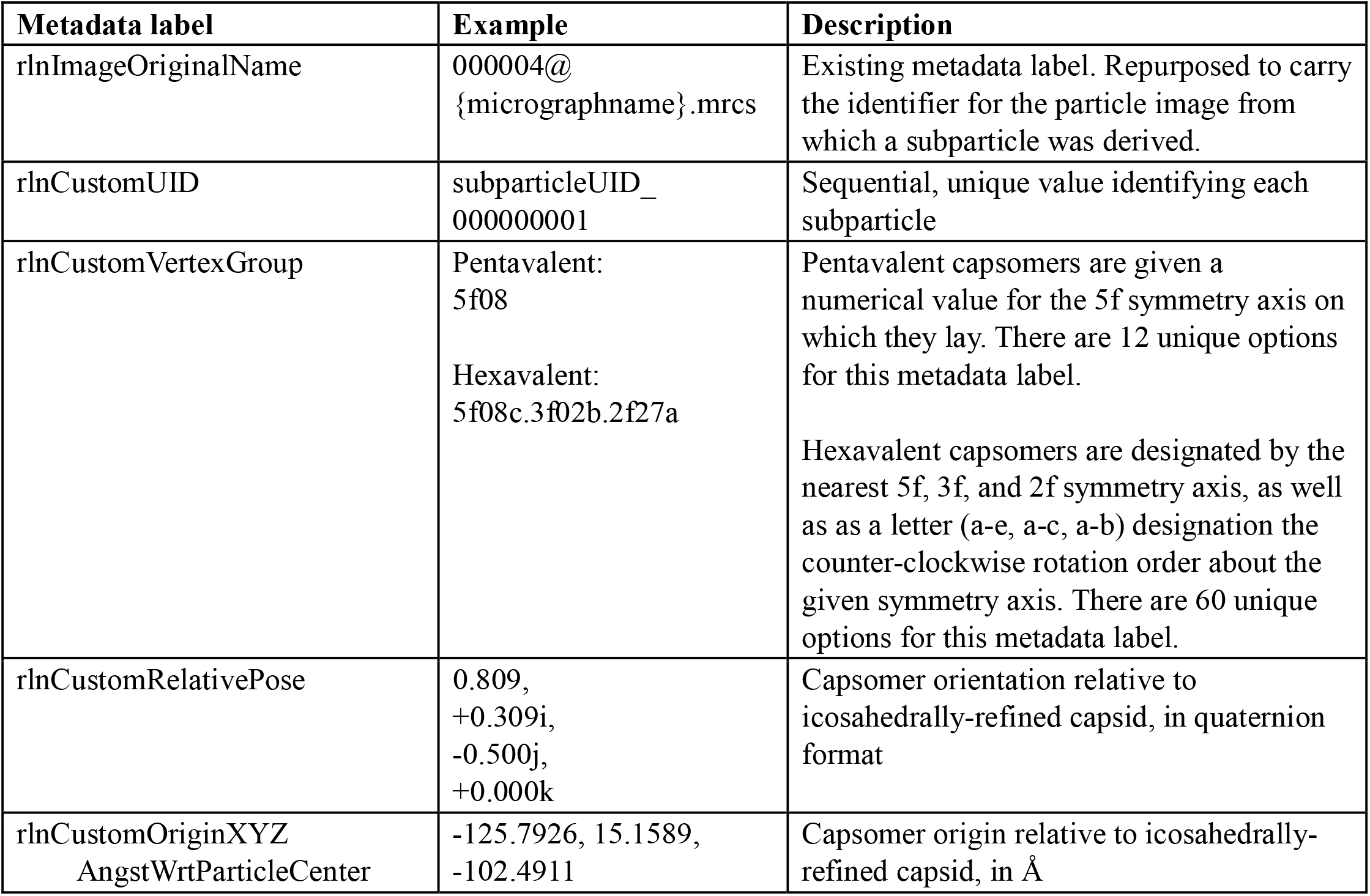
Custom ISECC metadata.

