## Supplemental Material for "High resolution cryo EM analysis of HPV16 identifies minor structural protein L2 and describes capsid flexibility"

### Supplementary Information

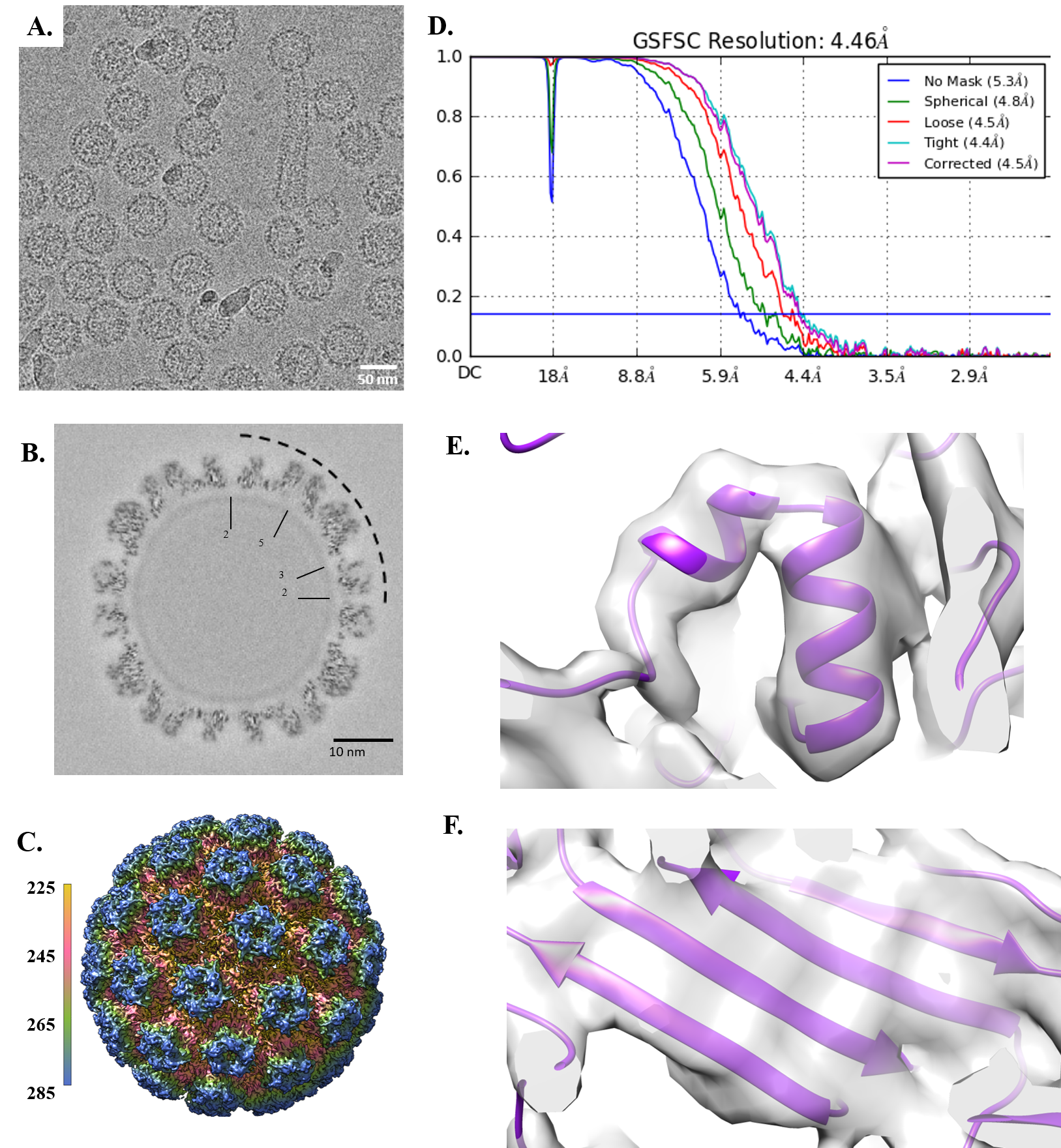

**SFig 1. Icosahedral Reconstruction.** (A) Representative micrograph 1,164 out of 8,936 collected. (B) Central cross-section of icosahedral structure with the symmetry labeled in the dotted region. (C) Full icosahedral reconstruction colored radially in Angstroms. (D) FSC Curve for icosahedral refinement. (E) Representative alpha helix (amino acid residues: 385-394, 396-401) at 4.5Å. (F) Representative beta sheet (amino acid residues: 71-76, 335-325, 153-160, 254-248) at 4.5Å.

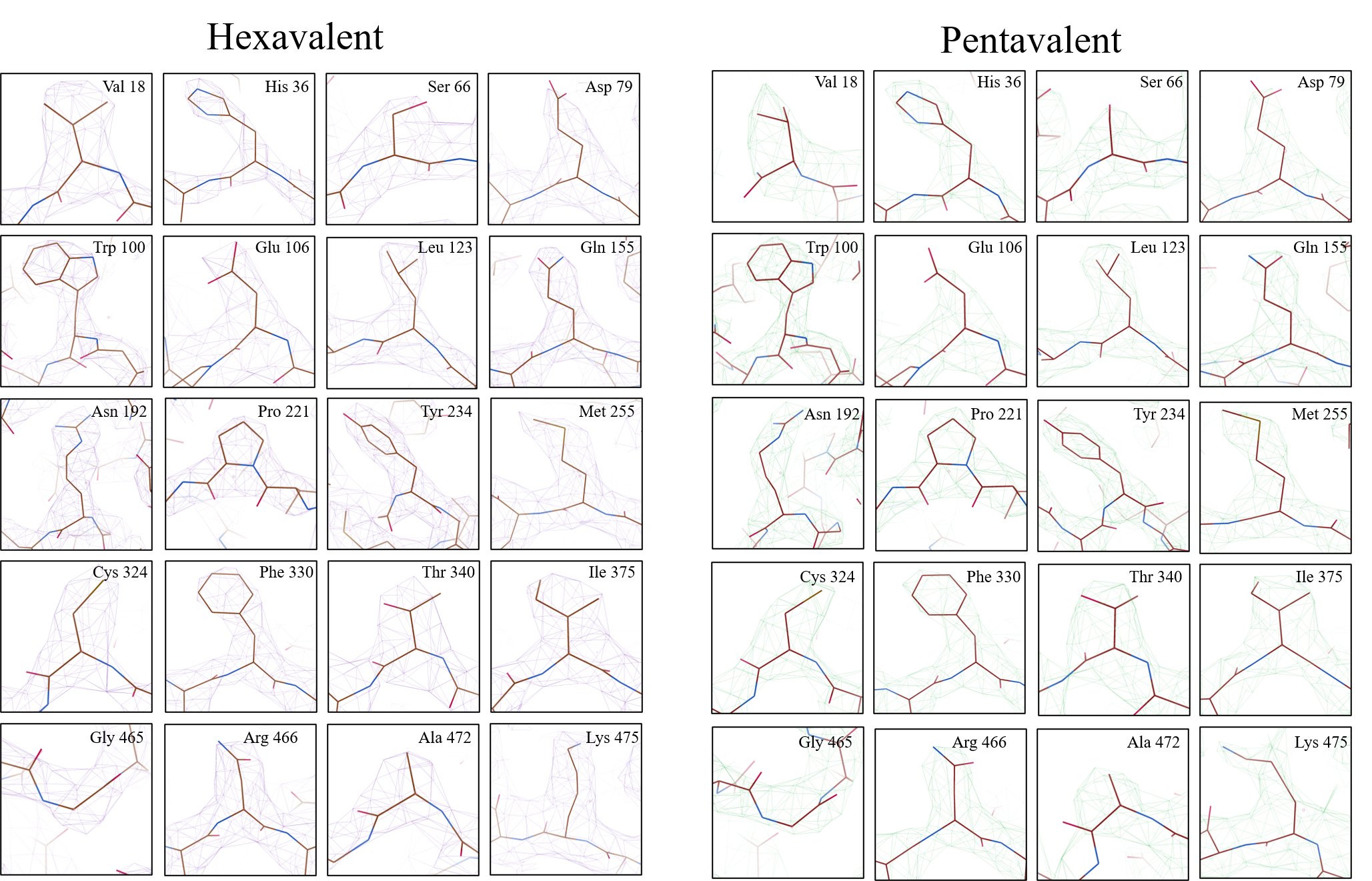

**SFig 2. Representative density of all side chains in pentavalent and hexavalent capsomers.** Representative side chains were chosen from L1 (brown) and compared between the pentavalent (purple density) and hexavalent (green density) capsomer environments.

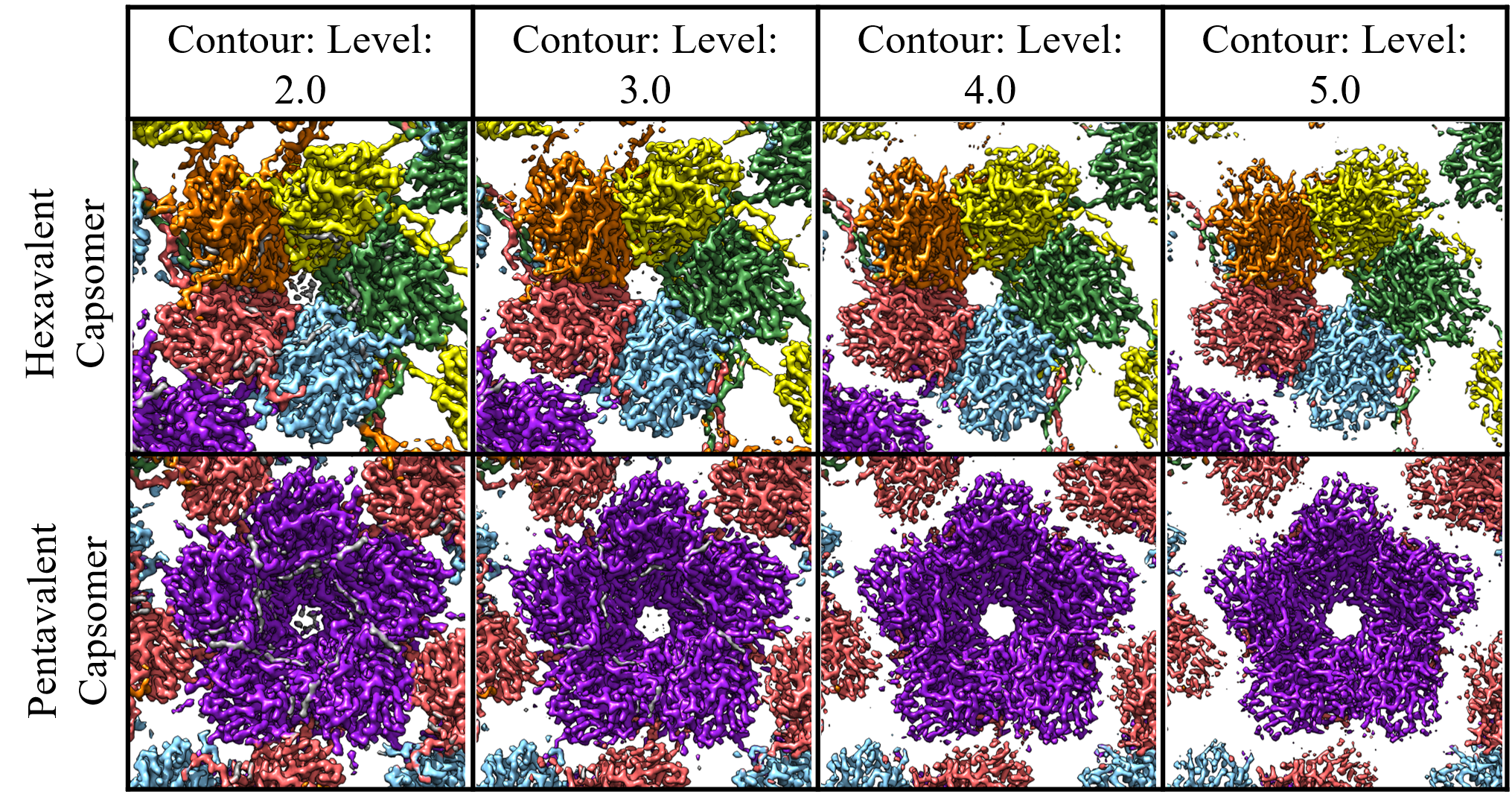

**SFig 3. L2 Density.** The surface rendered hexavalent (top) and pentavalent (bottom) capsomer density maps were colored as in Fig. 1A within 2Å of the L1 protein chain with unfilled density (gray) corresponding to L2. The L2 density is stronger in the pentavalent capsomer as can be seen with the changing contour of the map. In the hexavalent capsomers the same internal density can be noted, but the density disappears along with noise. Two distinct areas of gray density can be seen that are correlated to each chain of L1 in the capsomer.

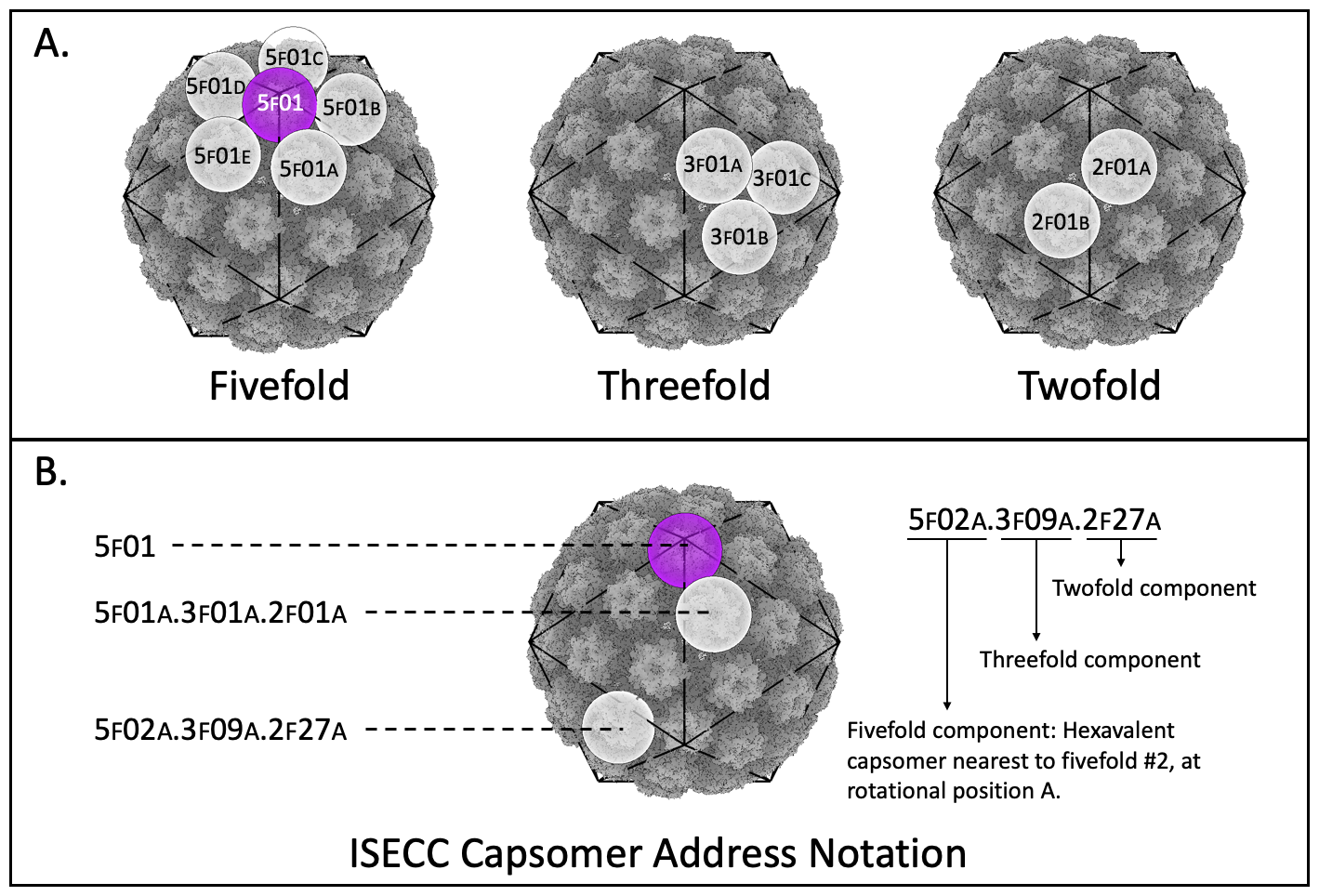

**SFig4. Capsomer addresses assigned in ISECC.** A) Each capsomer is assigned an address identifying the nearest symmetry vertices. Pentavalent capsomers only have a fivefold designation without any rotational parameter (left, purple). Hexavalent capsomers receive designations for the nearest fivefold (left), threefold (center), and twofold (right) axis, as well as rotational parameters. B) Examples of complete addresses implemented in ISECC are shown as one-part (pentavalent) and three-part (hexavalent) capsomer designations. Addresses are assigned during subparticle generation after normalization of the input vectors to a standard, shared asymmetric unit. This allows refinement parameters for any given subparticle to be correlated with other subparticles from the same parental particle.

**Supp. Table 1. Cryo-EM data collection, refinement and validation statistics**

| **Data Collection and Processing** | | **Icosahedral** | **Pentavalent Capsomer** | **Hexavalent Capsomer** |
| --- | --- | --- | --- | --- |
| Magnification | | 59,000 | 59,000 | 59,000 |
| Voltage (kV) | | 300 | 300 | 300 |
| Electron Exposure (e-/Å^2^) | | 60 | 60 | 60 |
| Defocus Range (um) | | 0.5-3.0 | 0.5-3.0 | 0.5-3.0 |
| Pixel Size (Å) | | 1.1 | 1.1 | 1.1 |
| Symmetry Imposed | | I1 | C5 | C1 |
| Micrographs Collected | | 10,143 | - | - |
| Micrographs Rejected (Bad Ice) | | 1,207 | - | - |
| Micrographs Accepted | | 8,936 | - | - |
| Initial Particle Number | | 202,705 | - | - |
| Final Particle Number | | 181,299 | 181,299 | 181,299 |
| Subparticles per Particle | | - | 12 | 60 |
| Final Subparticle Number | | - | 2,175,588 | 10,877,940 |
| Map Resolution (Å) | | 4.46 | 3.15 | 3.08 |
|  | FSC Threshold | 0.143 | 0.143 | 0.143 |
| **Refinement** | | **Recombined Icosahedral Asymmetric Unit** | | |
| Model composition | |  |  |  |
|  | Non-hydrogen atoms | 22492 | | |
|  | Protein Residues | 2864 | | |
| B-Factors | |  |  |  |
|  | Protein | - | | |
| R.m.s. Deviations | |  |  |  |
|  | Bond Length (Å) | 0.006 | | |
|  | Bond Angles (°) | 1.025 | | |
| Validation | |  |  |  |
|  | MolProbity Score | 2.65 | | |
|  | Clash Score | 13.17 | | |
|  | Rotamer Outliers (%) | 5.10 | | |
| Ramachandran Plot | |  |  |  |
|  | Favored (%) | 92.18 | | |
|  | Outliers (%) | 0.46 | | |

**Supplemental Table 2. Sequence Alignment of Ser306 – Ile328 Loop Region**

| **HPV Type** | **Overall Sequence Percent Identity** | **Sequence Percent Identity of Loop** | **Sequence** |
| --- | --- | --- | --- |
| 16 | - | - | SDAQIFNKPYWLQRAQGHNNGI |
| 31 | 82.97% | 92.3% | SDAQIFNKPYWMQRAQGHNNGI |
| 52 | 76.82% | 100% | SESQLFNKPYWLQRAQGHNNGI |
| 58 | 76.23% | 100% | SESQLFNKPYWLQRAQGHNNGI |
| 33 | 79.60% | 100% | SESQLFNKPYWLQRAQGHNNGI |
| 11 | 68.83% | 92.3% | SEAQLFNKPYWLQKAQGHNNGI |
| 6 | 68.59% | 92.3% | SEAQLFNKPYWLQKAQGHNNGI |
| 45 | 65.51% | 84.6% | SDSQLFNKPYWLHKAQGHNNGI |
| 18 | 65.87% | 84.6% | SDSQLFNKPYWLHKAQGHNNGV |

**Supplemental Table 3: Custom ISECC metadata**

| **Metadata label** | **Example** | **Description** |
| --- | --- | --- |
| rlnImageOriginalName | 000004@ {micrographname}.mrcs | Existing metadata label. Repurposed to carry the identifier for the particle image from which a subparticle was derived. |
| rlnCustomUID | subparticleUID_ 000000001 | Sequential, unique value identifying each subparticle |
| rlnCustomVertexGroup | Pentavalent:  5f08  Hexavalent: 5f08c.3f02b.2f27a | Pentavalent capsomers are given a numerical value for the 5f symmetry axis on which they lay. There are 12 unique options for this metadata label.   Hexavalent capsomers are designated by the nearest 5f, 3f, and 2f symmetry axis, as well as as a letter (a-e, a-c, a-b) designation the counter-clockwise rotation order about the given symmetry axis. There are 60 unique options for this metadata label. |
| rlnCustomRelativePose | 0.809, +0.309i, -0.500j, +0.000k | Capsomer orientation relative to icosahedrally-refined capsid, in quaternion format |
| rlnCustomOriginXYZ  AngstWrtParticleCenter | -125.7926, 15.1589,  -102.4911 | Capsomer origin relative to icosahedrally-refined capsid, in Å |
